# Direct Measurement of 8OG *syn-anti* Flips in Mutagenic 8OG•A and Long-Range Damage-Dependent Hoogsteen Breathing Dynamics Using ^1^H CEST NMR

**DOI:** 10.1101/2024.01.15.575532

**Authors:** Stephanie Gu, Hashim M. Al-Hashimi

## Abstract

Elucidating how damage impacts DNA dynamics is essential for understanding the mechanisms of damage recognition and repair. Many DNA lesions alter the propensities to form lowly-populated and short-lived conformational states. However, NMR methods to measure these dynamics require isotopic enrichment, which is difficult for damaged nucleotides. Here, we demonstrate the utility of the ^1^H chemical exchange saturation transfer (CEST) NMR experiment in measuring the dynamics of oxidatively damaged 8-oxoguanine (8OG) in the mutagenic 8OG*_syn_*•A*_anti_* mismatch. Using 8OG-H7 as an NMR probe of the damaged base, we directly measured 8OG *syn-anti* flips to form a lowly-populated (pop. ∼ 5%) and short-lived (lifetime ∼ 50 ms) non-mutagenic 8OG*_anti_*•A*_anti_*. These exchange parameters were in quantitative agreement with values from ^13^C off-resonance *R*_1ρ_ and CEST on a labeled partner adenine. The Watson-Crick-like 8OG*_syn_*•A*_anti_* mismatch also rescued the kinetics of Hoogsteen motions at distance A-T base pairs, which the G•A mismatch had slowed down. The results lend further support for 8OG*_anti_*•A*_anti_* as a minor conformational state of 8OG•A, reveal that 8OG damage can impact Hoogsteen dynamics at a distance, and demonstrate the utility of ^1^H CEST for measuring damage-dependent dynamics in unlabeled DNA.

## INTRODUCTION

DNA is continuously subjected to chemical damage, resulting in dozens of different chemical lesions which are repaired by specific damage repair enzymes^1-3^. The structural and dynamic properties of damaged DNA are of great interest not only because damage repair enzymes find and specifically bind to damaged DNA in a sea of excess undamaged DNA but also because damage repair frequently requires that the DNA cycle through alternative conformational states, which form as intermediates during multi-step catalytic cycles^4-13^. The dynamic properties of damaged DNA can vary with sequence context^14-20^, which could contribute to the sequence-dependence of damage repair frequently observed in many repair enzymes^14, 21^.

Many physiologically important motions in DNA involve non-native conformational states of base pairs (bps) that are typically lowly-populated and short-lived in naked, unbound DNA duplexes^16, 22-34^. Thus, it is highly desirable to have methods capable of efficiently assessing how damage impacts these micro-to-millisecond DNA dynamics. Solution NMR spectroscopy is one of the few experimental techniques capable of measuring the dynamics involving such fleeting states^35-39^. However, approaches such as spin relaxation in the rotating frame (*R*_1π_) and chemical exchange saturation transfer (CEST) require ^13^C and ^15^N isotopic enrichment of the DNA, which can be impractical and cost-prohibitive for damaged nucleotides. As a result, these NMR approaches that require isotopic enrichment are also ill-suited for broad and systematic explorations of how bp dynamics changes throughout the gamut of sequence, damage, and structural contexts. Thus, there is a need for more economic and efficient approaches that also bypass the requirement for isotopic enrichment.

Recently, we developed the imino ^1^H CEST experiment to assess the Watson-Crick to Hoogsteen exchange in unlabeled DNA samples^25^. Building on the SELective Optimized Proton Experiment^40^, the ^1^H CEST experiment targets the imino protons of nucleic acids to measure bp dynamics without the need for ^13^C/^15^N isotopic enrichment. Because ^1^H CEST data can be measured simultaneously for multiple imino resonances in a single experiment, the approach is ∼10-fold faster relative to conventional ^13^C/^15^N *R*_1π_ or CEST experiments employing Hartman-Hahn selective excitation ^41-42^. By eliminating the need for isotopic enrichment, the ^1^H CEST experiment could be used to broadly examine how damage shapes DNA dynamics. Here, using ^1^H CEST and the H7 imino proton of the oxidatively damaged 8-oxoguanine (8OG) and other imino resonances in Watson-Crick bps, we measured the dynamics of the mutagenic 8OG•A and its impact on the dynamics of neighboring bps within the duplex without isotopic enrichment of the DNA.

The 8OG lesion is one of the most common form of oxidative damage in DNA, occurring at least 1,000 times per day per cell if left unrepaired^43^. Whereas the undamaged G•A mismatch adopts G*_anti_*•A*_anti_* as the dominant conformation, the mutagenic 8OG lesion increases its propensity to form the 8OG*_syn_*•A*_anti_*Hoogsteen conformation^27, 44^. Because 8OG*_syn_*•A*_anti_* mimics the Watson-Crick geometry (Fig. 1A), it can evade DNA polymerase fidelity checkpoints, increasing the frequency of replication errors and leading to mutations linked to colorectal, lung, and esophageal cancers^45-47^.

**Figure 1.**
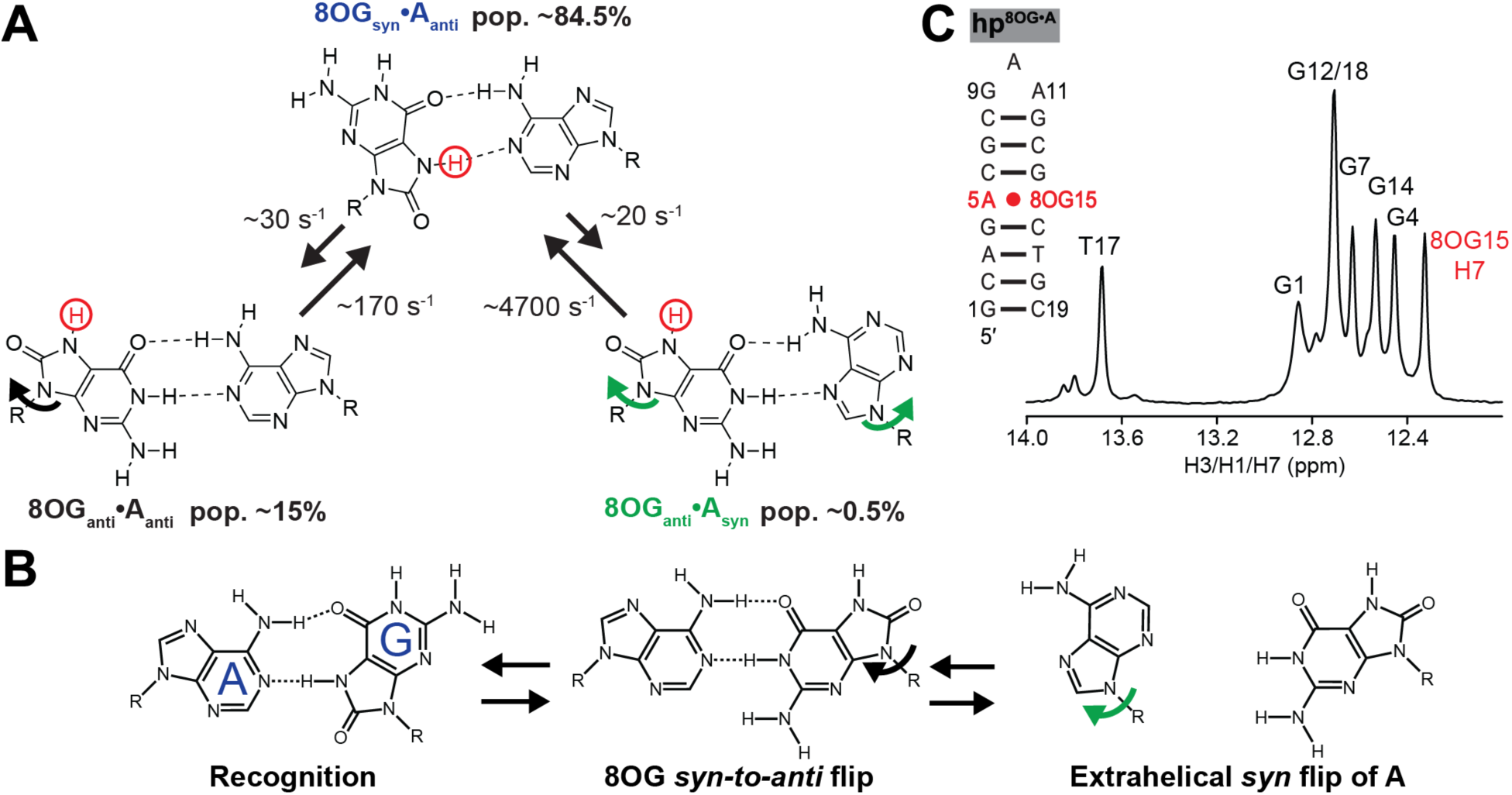
Measuring the dynamics of the 8OG•A mismatch using ^1^H CEST. (A) Dynamic equilibrium previously determined for 8OG•A using ^13^C *R*_1π_ and CEST NMR experiments targeting the adenine residue^27^. The dominant 8OG*_syn_*•A*_anti_*ground-state conformation exists in dynamic equilibrium with two excited conformational states: 8OG*_anti_*•A*_anti_* forms by flipping the 8OG base from *syn* to *anti* and 8OG*_anti_*•A*_syn_*forms through dual flipping of both the 8OG and A bases. The 8OG-H7 imino proton used in ^1^H CEST experiments is highlighted in red. (B) Proposed kinetic mechanism for A•8OG damage repair by MutY/MUTYH involves the excited conformational states as intermediates^14, 27^. (C) The DNA hairpin (hp^8OG•A^) used to study the dynamics of 8OG•A by NMR. Also shown is the 1D ^1^H spectrum of the imino region at T = 10°C highlighting the 8OG-H7 imino resonance in red. The buffer was 25 mM sodium chloride, 15 mM sodium phosphate pH 7.4, 0.1 mM EDTA, and 10% D_2_O.

Recently, an NMR study showed that the 8OG•A mismatch is not locked into the mutagenic 8OG*_syn_*•A*_anti_*ground state (GS) but can transiently form two non-mutagenic excited state conformational states (ESs) 8OG*_anti_*•A*_anti_* and 8OG*_anti_*•A*_syn_* through *syn-anti* flips of the purine bases^27^ (Fig. 1A). The 8OG*_anti_*•A*_anti_* ES had a comparatively high population (pop.) of ∼15% and slow *k*_ex_ = *k*_forward_ + *k*_backward_ ∼ 200 s^-1^. It forms by flipping the 8OG purine base about its glycosidic bond from the *syn* to the *anti* conformation (Fig. 1A). The 8OG*_anti_*•A*_syn_* with a lower pop. of ∼0.5% and faster k_ex_ ∼ 4,700 s^-1^ was proposed to form through dual flips of the two purine bases (Fig. 1A).

In both cases, the flipping of 8OG could only be indirectly inferred based on NMR chemical exchange measurements on the partner adenine-C8/C1ʹ and chemical shift fingerprinting of conformational mimics^27^.

The propensities to form non-mutagenic 8OG*_anti_*•A*_anti_* and 8OG*_anti_*•A*_syn_* could vary with sequence, thereby altering the mutagenicity of 8OG. This could in turn give rise to sequence-dependent replication errors linked to cancer mutational signatures^46-48^. The conformational dynamics of 8OG•A is also important for understanding its repair mechanism. Both 8OG*_anti_*•A*_anti_* and 8OG*_anti_*•A*_syn_* are proposed to form as intermediates during 8OG repair^4-10, 13, 27, 49-55^. Starting with the 8OG*_syn_*•A*_anti_* initially recognized by the base excision repair enzyme MutY/MUTYH, the 8OG base flips into the *anti* conformation to form 8OG*_anti_*•A*_anti_*^6^ (Fig. 1B). The adenine base then flips into the *syn* conformation to form 8OG*_anti_*•A*_syn_* followed by extra-helical flipping so that the base occupies the MutY/MUTYH active site for cleavage^12^ (Fig. 1B). Again, the dependence of these 8OG•A flipping dynamics on sequence could potentially contribute to the observed sequence-dependence of 8OG damage and its repair linked to cancer mutational signatures^46-48^. Finally, it has also been proposed that 8OG can act as an epigenetic-like marker regulating transcription where its flexibility and propensity to form alternative conformation in the DNA double helix could influence transcription factor binding and subsequent gene expression^56^. Thus, it is of great interest to measure the dynamics of the 8OG•A bp, its sequence dependence, and its potential to impact the dynamics of neighboring bps and the global overall DNA.

Here, we demonstrate the utility of the ^1^H CEST experiment in directly measuring *syn-anti* flips of the damaged 8OG base as well as the impact of 8OG on the dynamics of other bps without the need for isotopic enrichment of the DNA. At two different temperatures, the exchange parameters measured by ^1^H CEST for 8OG*_anti_*•A*_anti_* were in quantitative agreement with values measured using ^13^C off-resonance *R*_1π_ and CEST experiments on the ^13^C/^15^N isotopically labeled partner adenine. The results support the identity of the non-mutagenic 8OG*_anti_*•A*_anti_* as a minor conformational state of the 8OG•A mismatch ensemble, reveal how 8OG damage can affect Hoogsteen breathing at distant Watson-Crick bps, as well as demonstrate the utility of ^1^H CEST for measuring damage-dependent DNA dynamics. With ^1^H CEST, it should be possible to comprehensively measure the sequence-dependence of 8OG•A dynamics in the future.

## Experimental Methods

### Sample Preparation

#### Unlabeled DNA oligonucleotides

Unlabeled DNA oligonucleotides (hp^A-T^ and hp^G•A^) were purchased from Integrated DNA Technologies with standard desalting purification. The unlabeled modified DNA oligonucleotide contains 8OG (hp^8OG•A^) was purchased from the Yale Keck Oligonucleotide Synthesis Facility with cartridge purification using commercially sourced amidites from Glen Research and synthesized on a Dr. Oligo 192 or an Applied Biosystems 394 synthesizer.

#### 13C,15N-labeled DNA samples

Uniformly ^13^C,^15^N-labeled hp^8OG•A^ was purchased from and synthesized by the Yale Keck Oligonucleotide Synthesis Facility as described previously^27^.

#### NMR buffer

The NMR experiments employed a sodium phosphate buffer prepared by the addition of equimolar solutions of sodium phosphate monobasic and dibasic salts, sodium chloride, and EDTA to give final concentrations of 15 mM sodium phosphate, 25 mM sodium chloride and 0.1 mM EDTA. The pH was adjusted to 7.4 by adding phosphoric acid or sodium hydroxide. The buffers were then brought up to the desired volume, vacuum filtered, and stored for usage.

#### Sample annealing and buffer exchange

Oligonucleotides were resuspended in water and annealed by heating at a temperature of 95°C for ∼5 min, followed by cooling on ice for ∼1 h. Samples were then exchanged into the NMR buffer using Amicon Ultra-4 centrifugal concentrators (4 mL; Millipore Sigma) with a 3-kDa molecular cutoff to a final volume of ∼250 μL. Deuterium oxide (10% vol/vol) was added to the samples before the NMR experiments.

### NMR Spectroscopy

NMR experiments were performed on 600, 700, and 900 MHz Bruker Avance spectrometers equipped with HCNP, HCN, and HCN cryogenic probes, respectively, running TopSpin 3.2 (600 and 700 MHz) or TopSpin 4 (900 MHz). All experiments were performed in NMR buffer at a temperature of 25°C unless stated otherwise. The NMR data were processed and analyzed using NMRPipe^57^ and SPARKY (T. D. Goddard and D. G. Kneller, SPARKY 3, University of California, San Francisco), respectively.

#### Nuclear magnetic resonance assignments

Sugar and base resonances of hp^A-T^ were assigned using 2D [^1^H,^1^H] NOESY and 2D[^13^C,^1^H] HSQC experiments measured at pH 7.4 and 25°C as described previously^27^. The resonance assignments for hp^G•A^ and hp^8OG•A^ were reported previously^27^.

### 1H CEST

The ^1^H CEST experiments were performed using a pulse sequence described recently^25^. The spin-lock powers used ranged from 10 to 4000 Hz (Table S1). The relaxation times (50 ms or 100 ms) were chosen to achieve a ∼40-50% loss in signal intensity when comparing the resonance intensity without delay versus with delay at a far off-resonance offset with a spin-lock power of 10 Hz. In the case of 8OG-H7, a relatively short 50 ms delay time was also chosen to minimize NOE effects^25^. Calibration of radiofrequency (RF) fields was performed as described previously^25^. The experimental conditions for all ^1^H CEST experiments (temperature and magnetic field strength) are summarized in Table S1.

#### Fitting of ^1^H CEST data

One-dimensional peak intensities were calculated using NMRPipe^57^. The error in the intensity for a given spin-lock power was determined based on the standard deviation from triplicate (n = 3) measurements with a zero relaxation delay using the same spin-lock power. The intensities were normalized to the average intensity of the three zero relaxation delay measurements. The CEST profiles were generated by plotting the normalized intensity as a function of offset Θ_OBS_ = χο_RF_-χο_OBS_ in which χο_OBS_ is the Larmor frequency of the observed resonance, and χο_RF_ is the angular frequency of the applied RF field. The RF field inhomogeneity was accounted for during the fitting as described previously^25^. The data was fit with and without 2-state exchange by fitting the intensity values to the numerical solution of the Bloch-McConnell (B-M) equations^36^. The reduced chi-square (rχ^2^) was calculated to assess the goodness of fit. Model selection for the fits with and without exchange were evaluated by computing the Akaike’s (AIC)^58^ and Bayesian information criterion (BIC)^59^ weights. The improvement in the fit was considered statistically significant if both wAIC_+ex_ and wBIC_+ex_ values were > 0.995 and rχ^2^ was reduced with the inclusion of exchange. The fitted exchange parameters which include the population of the ES (*p*ES), the exchange rate between the GS and the ES (*k*ex = *k*forward + *k*reverse), and the chemical shift difference between the ES and GS conformations (Δω = χο_ES_-χο_GS_ in which χο_ES_ and χο_GS_ are the chemical shifts of the ES and GS, respectively) are summarized in Tables S2 and S4. The errors in the exchange parameters were determined based on the fitting errors which were obtained as the square root of the diagonal elements of the covariance matrix.

#### *Off-resonance ^13^C R*_1π_ relaxation dispersion

*^13^C R*_1π_ experiments were performed on the adenine-C1ʹ or adenine-C8 of hp^8OG•A^ at 10°C in NMR buffer ^42^. The magnetization corresponding to adenine-C1ʹ or adenine-C8 was allowed to relax under an applied spin-lock field for a maximal duration (60 ms for adenine-C1ʹ and 44 ms for adenine-C8) chosen to achieve ∼ 70% loss in signal intensity at the end of the relaxation period. The signal intensity was recorded for five delays equally spaced over the relaxation period. Spin-lock powers used for the *R*_1π_ measurements ranged between 100 and 3,000 Hz. Absolute offset frequencies were chosen ranging from zero to 3.5 times the given spin-lock power. Briefly, the *R*_1π_ experiment measures the line-broadening contribution (*R*_ex_) to the transverse relaxation rate (*R*_2_) during a relaxation period where a continuous RF field is applied with variable power (μ_SL_) and frequency (μ_RF_). For a system experiencing detectable exchange, the relaxation dispersion (RD) profiles depicting the dependence of *R*_2_ + *R*_ex_ on μ_SL_ and μ_RF_ show a characteristic peak typically centered about the difference between the chemical shift of the ground state (GS) and the excited state (ES) (Δm = μ_ES_-μ_GS_). The experimental conditions for ^13^C *R*_1π_ experiments including the magnetic field, spin-lock powers, and offsets used are listed in Table S1.

#### Fitting of ^13^C R_1ρ_ data

The ^13^C *R*_1π_ values for a given spin-lock power and offset combination were obtained by fitting the peak intensities (extracted using NMRPipe^57^) as a function of delay time to a mono-exponential decay function. The B-M equations^36^ were used to fit *R*_1π_ values for either adenine-C1ʹ or adenine-C8 as a function of spin-lock power and offset to a 2-state exchange model assuming *R*2,GS = *R*2,ES and *R*_1,GS_ = *R*_1,ES_^35-36^. Like the fitting of the ^1^H CEST data, the fitting of the ^13^C *R*_1π_ data yielded exchange parameters of interest, including the population of the ES (*p*ES), the exchange rate between the GS and the ES (*k*ex = *k*forward + *k*reverse), and the chemical shift difference between the ES and GS conformations (Δω = χο_ES_-χο_GS_ in which χο_ES_ and χο_GS_ are the chemical shifts of the ES and GS, respectively) with the uncertainties in the exchange parameters determined using a Monte Carlo procedure^35^. The fitted parameters are summarized in Table S3.

Off-resonance *R*_1π_ profiles were generated by plotting (*R*_2_ + *R*_ex_) = (*R*_1π_-*R*_1_cos^2^8)/sin^2^8 where 8 is the angle between the effective field of the observed resonance and the z-axis in radians as a function of Ο_eff_ = χο_OBS_ - χο_RF_ where χο_OBS_ is the Larmor frequency of the observed resonance, and χο_RF_ is the carrier frequency of the applied spin-lock power. The uncertainty in (*R*_2_ + *R*_ex_) was determined by propagating the error in *R*_1π_ through a Monte Carlo procedure.

## Results and Discussion

### H7 is a direct NMR probe of 8OG *syn-anti* flips

8OG has a unique H7 imino proton, which is hydrogen-bonded in the dominant 8OG*_syn_*•A*_anti_* GS conformation (Fig. 1A). The 8OG-H7 resonance has an NMR chemical shift (∼12.4 ppm)^27, 60^ like that of G-H1 in a canonical Watson-Crick bp (∼12-13 ppm)^25^. When the 8OG base flips to the *anti* conformation in 8OG*_anti_*•A*_anti_* and 8OG*_anti_*•A*_syn_*, the 8OG-H7 imino is no longer hydrogen-bonded. The G-H1 undergoes a similar transition in hydrogen-bonding when it flips from the *anti* conformation in the Watson-Crick G-C bp to the *syn* conformation in the G*_syn_*-C^+^ Hoogsteen bp, and this is accompanied by a sizeable upfield shift in the G-H1 of ∼2-3 ppm^25^. Thus, we can expect a similar upfield shift for 8OG-H7 when 8OG adopts the *anti* conformation, making it a potentially powerful ^1^H CEST probe of 8OG base flipping. While in theory ^1^H CEST could also be used to measure the dual flips of 8OG and A to form 8OG*_anti_*•A*_syn_*, simulations show that the 8OG*_anti_*•A*_syn_* exchange contribution will be very small due to its lower population and faster exchange kinetics (Fig. S1).

### Resolving 8OG*_anti_*•A*_anti_*

We used ^1^H CEST to measure conformational exchange in an unlabeled DNA hairpin hp^8OG•A^ (∼2 mM) containing a central 8OG•A mismatch (Fig. 1C). We previously showed^27^ that the mismatch forms 8OG*_syn_*•A*_anti_*as the dominant GS conformation. The ^1^H CEST experiments were performed at pH 7.4 and T = 10°C unless indicated otherwise. Data was collected using a 50 ms relaxation delay and required ∼6 hours on 900 MHz Bruker NMR spectrometers equipped with cryogenic probes (See Experimental Methods). At low temperature, the 8OG-H7 resonance was well-resolved and relatively isolated from other imino resonances (Fig. 1C) while the 8OG-H1 resonance was not observable as expected due to rapid solvent exchange^61^.

Briefly, the CEST experiment measures the impact of conformational exchange on longitudinal GS magnetization during a relaxation period in which a continuous RF field is applied with variable power (μ_SL_) and frequency (μ_RF_). When applied on-resonance with the chemical shift of an ES, the RF field saturates the ES magnetization, which can then be transferred via conformational exchange to the GS. This reduces the signal intensity for the GS, typically resulting in a minor dip centered about μ_ES_ = Δm when the RF is on-resonance with the ES chemical shift. Likewise, a major dip is also observed at μ_GS_ = 0. The CEST data can be fit to the B-M equations^36^ to determine the rate of exchange (*k*_ex_), population of the ES (pop.), and difference between the ES and GS chemical shifts (Δm = μ_ES_-μ_GS_).

When using the ^1^H CEST experiment, one must carefully consider the contributions from proton-proton cross-relaxation, which can lead to spurious Nuclear Overhauser Effect (NOE) dips in the ^1^H CEST profiles^25, 38, 40, 62^. These contributions were suppressed by selective excitation of the 8OG-H7 and dephasing of non-imino protons before applying the *B*_1_ field and by using a short relaxation delay of 50 ms as described previously^25^. In addition, in 2D [^1^H, ^1^H] NOESY spectra, we did not observe cross-peaks between 8OG-H7 and any resonance within the 4-ppm range at 25°C (Fig. S2). Upon reducing the temperature to 1°C, we observed a cross-peak ∼2.5 ppm upfield shifted from 8OG-H7, but this NOE cross-peak did not correspond to any dips in the ^1^H CEST profiles (Fig. S2). Although the adenine amino protons are in proximity to the partner 8OG-H7, these amino protons were not observable in 1D ^1^H or 2D [^1^H, ^1^H] NOESY spectra due to the intermediate exchange about the C-N bond of the amino.

We recently demonstrated the feasibility of applying high spin-lock powers in CEST experiments, broadening the timescales accessible to the experiment^25, 41^. We used a broad range of spin-lock powers spanning 10 to 1,000 Hz to measure the 8OG-H7 ^1^H CEST profiles. Indeed, at T = 10°C, we observed a clear minor dip in the ^1^H CEST profile of 8OG-H7 indicating conformation exchange with a lowly-populated and short-lived ES (Fig. 2). The dip was upfield shifted by Δm ∼ −2 ppm, which is in excellent agreement with the Δm ∼ −2 ppm experienced by G- H1 when transitioning between the G*_anti_*-C Watson-Crick and G*_syn_*-C^+^ Hoogsteen bps^25^. The dip also occurred at an offset frequency that does not correspond to any other observable proton frequency in hp^8OG•A^, and thus is unlikely to be the result of NOE effects (Fig. S2). Based on Akaike’s (AIC) and Bayesian information criterion (BIC) analysis^58-59^, we obtained a statistically significant improvement in fitting the 8OG-H7 ^1^H CEST profile when including 2-state but not 3- state chemical exchange (Fig. S3).

**Figure 2.**
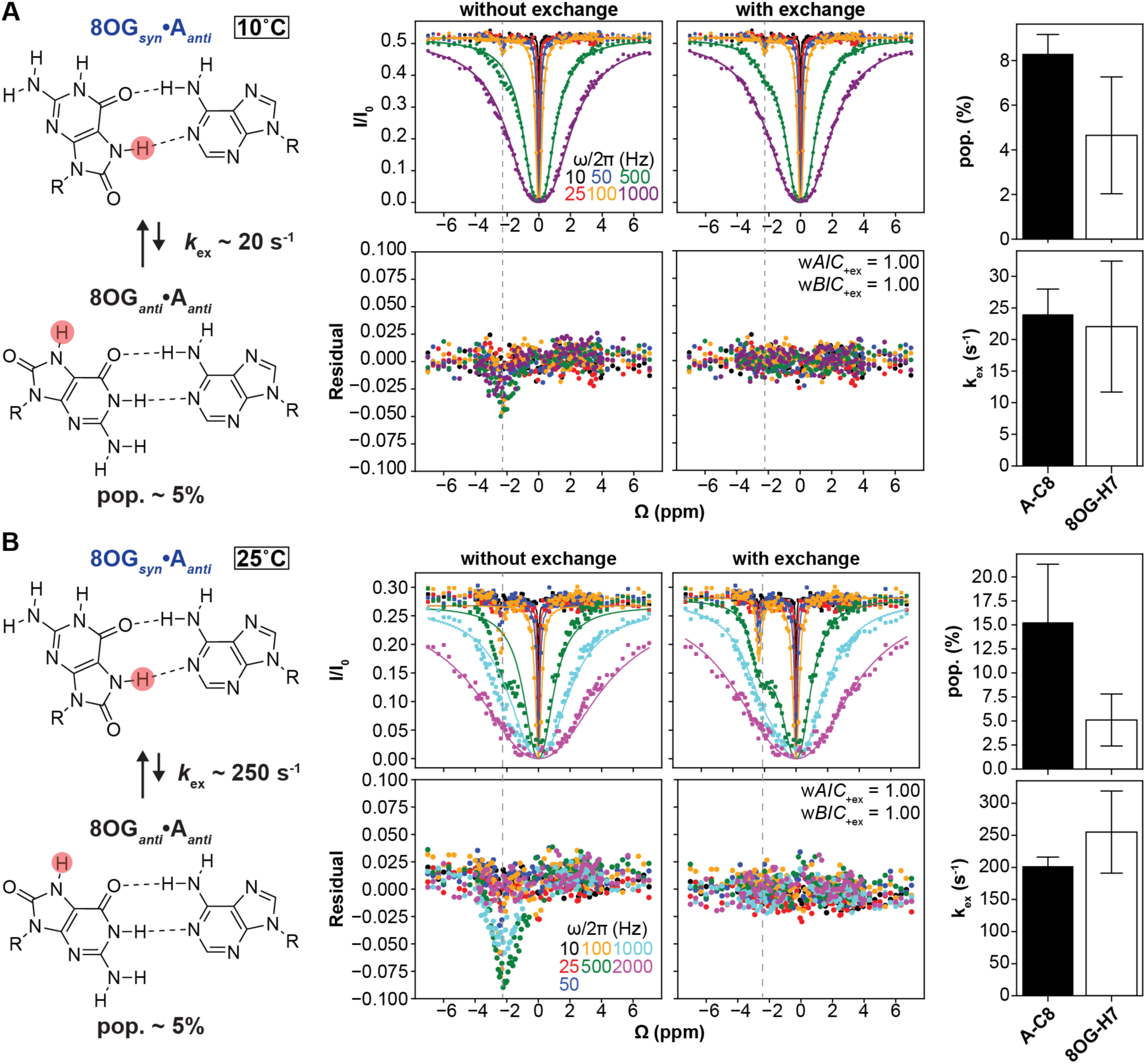
Resolving conformational exchange involving 8OG*_anti_*-A*_anti_*in hp^8OG•A^ using ^1^H CEST. (A) ^1^H CEST profiles for 8OG-H7 measured at (A) T = 10°C and (B) T = 25°C. Shown are the fits to the ^1^H CEST data using B-M equations^36^ with and without 2-state chemical exchange. Below the ^1^H CEST profiles are the residual plots (experimental normalized intensity – fitted normalized intensity). Also shown in inset are Akaike’s (*wAIC*) and Bayesian information criterion (*wBIC*) weights for fits with exchange. The dashed gray lines indicate the 8OG*_anti_*•A*_anti_* Δω position in both ^1^H CEST profiles and in the residual plots. The error bars for the ^1^H CEST profiles, which are smaller than the data points, were obtained using triplicate experiments. RF powers are color-coded. Also shown are the fitted exchange parameters measured by ^1^H CEST as well as ^13^C *R*_1π_ and CEST. Errors in exchange parameters were fitting errors of ^1^H CEST, which were calculated as the square root of the diagonal elements of the covariance matrix, and of ^13^C *R*_1π_ and CEST, which were calculated using a Monte-Carlo scheme^35^. The exchange parameters at 25°C for comparison to ^1^H CEST were determined from ^13^C CEST previously reported in Gu *et al^27^*.

A 2-state fit of the 8OG-H7 ^1^H CEST profile yielded an ES population of 5 ± 3% and exchange rate *k*_ex_ = 20 ± 10 s^-1^ with Δω_8OG-H7_ = −2.27 ± 0.01 ppm (Fig. 2A and Table S2) in excellent agreement with values measured for 8OG*_anti_*•A*_anti_*exchange using independent ^13^C off-resonance *R*_1π_ measurements on the adenine-C8 (Table S3). When we subjected the ^1^H CEST data to a degeneracy analysis, the rχ^2^ increased significantly when varying *k*_ex_, Δω, or *p*_ES_ by 3-fold or 1 ppm (Fig. S4A-B). The <2-fold difference between the populations obtained using ^1^H CEST and ^13^C *R*_1π_ are within error and comparable to those reported previously when comparing the exchange kinetics of Watson-Crick to Hoogsteen transition^25^.

To verify these results, we repeated the 8OG-H7 ^1^H CEST measurements at a higher temperature of 25°C using spin-lock powers from 10 to 2,000 Hz. At the higher temperature, the 8OG-H7 resonance was less well-resolved due to exchange broadening but still sufficiently isolated to be targetable by ^1^H CEST (Fig. S4C). Once again, we observed a minor dip at Δω ∼ 2 ppm, but it was substantially more intense compared to T = 10°C (Fig. 2B). A 2-state fit of the ^1^H CEST profiles yielded Δω_8OG-H7_ = −2.36 ± 0.01 ppm as expected for 8OG*_anti_*•A*_anti_* and like the Δω measured at T = 10°C. The exchange parameters *p*_ES_ = 5 ± 3% and exchange rate *k*_ex_ = 260 ± 60 s^-1^ (Fig. S4B-C and Table S2) were in good agreement with counterparts measured independently using ^13^C CEST measurements on the adenine-C8 as reported previously^27^ (Fig. S4C). The larger dip observed at the higher temperature could be attributed to the nearly 10-fold increase in *k*_ex_. Again, when we subjected the ^1^H CEST data to a degeneracy analysis, the rχ^2^ increased significantly when varying *k*_ex_, *p*_ES_, or Δm by 3-fold or 1 ppm (Fig. S4A). The ∼3-fold lower population in the ^1^H CEST data relative to the ^13^C CEST data at the higher temperature of 25°C could be attributed to the 8OG-H7 undergoing solvent exchange, partial overlap of the 8OG-H7 resonance with other imino resonances, or both. Together, these temperature-dependent data establish the utility of ^1^H CEST to directly measure the 8OG flip to form the minor 8OG*_anti_*•A*_anti_*conformational state.

Because of its low population and fast exchange kinetics, we were unable to quantitatively resolve an exchange contribution from 8OG*_anti_*•A*_syn_* especially given that the fast exchange is expected to create asymmetry in the major dip rather than manifest itself as a separate minor dip (Fig. S1). Moreover, any asymmetry in the major dip is likely to be masked by the larger contribution from 8OG*_anti_*•A*_anti_* (Fig. S1). Nevertheless, the ^1^H CEST profiles measured at spin-lock powers ≥ 1,000 Hz at T = 10°C, which are conditions predicted to optimally isolate the 8OG*_anti_*•A*_syn_* exchange contribution from 8OG*_anti_*•A*_anti_* rather than 8OG*_anti_*•A*_anti_* (Fig. S5).

### The G-A mismatch slows Hoogsteen breathing at distant Watson-Crick base pairs

One of the advantages of ^1^H CEST is the ability to probe multiple protons simultaneously, which allows for duplex-wide analysis of bp dynamics. Prior studies showed that Watson-Crick A-T and G-C bps exist in dynamic equilibrium with lowly-populated and short-lived Hoogsteen bps which form by flipping the purine base from the *anti* to the *syn* conformation^16, 22-23, 25, 29, 33-34, 63-64^. The Hoogsteen breathing motions have been implicated in protein-DNA recognition^22, 30, 65^, damage induction^29, 63^, and DNA replication^66^. Prior studies showed the utility of ^1^H CEST to measure these fundamental Hoogsteen breathing motions^25^. Thus, we used ^1^H CEST to examine how replacing the A-T bp at the center of the hairpin (hp^A-T^) with either G•A (hp^G•A^) or 8OG•A (hp^8OG•A^) (Fig. 3) impacts the Watson-Crick to Hoogsteen exchange dynamics of other bps in the DNA duplex.

**Figure 3.**
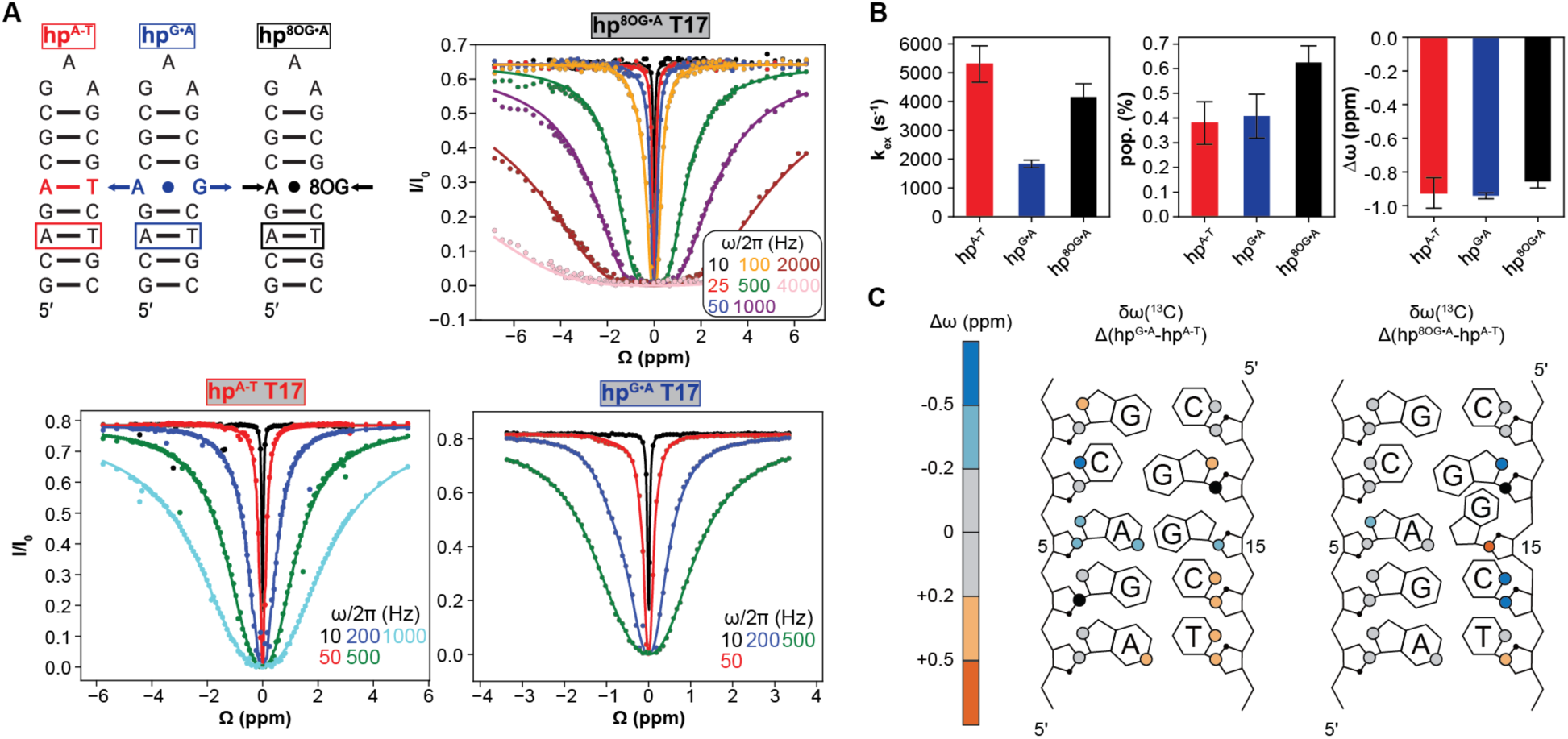
Impact of G•A and 8OG•A on the Hoogsteen breathing dynamics of other base pairs in the DNA duplex. (A) DNA hairpins used to compare dynamics upon modifying the central bp. The T17 base with detectable Watson-Crick to Hoogsteen exchange is highlighted in the box. Arrows at the mismatch are used to indicate expansion and constriction of the C1ʹ-C1ʹ distance for G•A (∼12 Å) and 8OG•A (∼10.5 Å), respectively. Shown are the T17-H3 ^1^H CEST profiles measured in hp^8OG•A^ (top right), hp^A-T^ (bottom left), and hp^G•A^ (bottom right) at 15°C and the fits to 2-state exchange. The error bars for the ^1^H CEST profiles, which are smaller than the data points, were obtained using triplicate experiments as described in Experimental Methods. RF powers are color-coded. (B) Comparison of fitted exchange parameters (pop., *k*ex, and Δw) across the three hairpins. (C) Comparison of ^13^C chemical shift perturbations (δω) induced by G•A (Δ(hp^G•A^-hp^A-T^)) versus 8OG•A (Δ(hp^8OG•A^-hp^A-T^)) relative to A-T for the indicated resonances. The οχο values are color-coded on the duplexes. The atoms colored in black could not be reliably assigned due to their proximity to their water resonance.

At pH = 7.4, it is normally feasible to measure Watson-Crick to Hoogsteen exchange contribution for A*_syn_*-T bps but not protonated G*_syn_*-C^+^ bps whose detection requires lower pH conditions^67^. Recently, it was shown that due to fast exchange kinetics, detection of Hoogsteen breathing even for A-T bps can be difficult when sandwiched by G-C neighbors (Manghrani *et al* submitted). Thus, we expected that the bps in the hp^A-T^ Watson-Crick reference would not experience any detectable Hoogsteen exchange under room temperature conditions.

Indeed, at T = 25°C, the ^1^H CEST profiles measured in hp^A-T^ for T17, T15, G18, G7, and G4 did not show any signs of exchange (Fig. S6). G12 and G14 could not be reliably measured due to spectral overlap. Upon lowering the temperature to T = 15°C (Fig. S7), the central T15 (pop. ∼ 0.06%, *k*_ex_ ∼ 3200 s^-1^, Δω ∼ −1.9 ppm) and T17 (pop. ∼ 0.4%, *k*_ex_ ∼ 5300 s^-1^, Δω ∼ −0.9 ppm) showed exchange consistent with an A*_syn_*-T Hoogsteen ES^25^ (Fig. 3A and Table S4). No exchange was detected at G-C bps, as expected. The absence of exchange at T = 25°C for the A-T bps was most likely because the Hoogsteen exchange was too fast, falling outside detection.

For hp^G•A^ in which the central A-T bp was replaced with a G•A mismatch, the ^1^H CEST profiles measured at T = 15°C showed no sign of exchange for all G-C bps. Based on the G15-H1 CEST profile measured for the central G•A bp, we observed the expected Hoogsteen exchange to form G*_anti_*•A*_syn_* as reported previously^27^. Interestingly, we measured an exchange contribution for T17- H3, and a 2-state fit of the ^1^H CEST profile yielded a similar population (pop. ∼ 0.5%) but ∼2-fold slower exchange rate (*k*_ex_ ∼ 1800 s^-1^) relative to that measured for the same bp in the Watson-Crick hp^A-T^ control (Fig. 3). This ∼2-fold reduction in the exchange rate was statistically significant as fixing the exchange parameters to those measured in hp^A-T^ resulted in a much poorer fit to the ^1^H CEST profile (Fig. S8). This ∼2-fold slowdown in Hoogsteen breathing could be the consequence of the G*_anti_*•A*_anti_* conformation having a larger C1ʹ-C1ʹ distance (∼12 Å) that disfavors the formation of the more constricted (∼8.5 Å) A-T Hoogsteen bp. Indeed, T-T mismatches, with constricted C1ʹ-C1ʹ distance (∼9 Å), were previously shown to promote Hoogsteen breathing^22, 28^. Thus, the replacement of A-T with G•A led to a measurable impact on the Hoogsteen dynamics two bps away.

### The 8OG lesion restores Hoogsteen breathing at distant Watson-Crick base pairs

Because 8OG*syn*•A*anti* better mimics the Watson-Crick-like C1ʹ-C1ʹ distance (∼10.5 Å) relative to G*anti*•A*anti*, one might expect that the Hoogsteen dynamics measured in damaged hp^8OG•A^ would more closely resemble those of the Watson-Crick hp^A-T^ control. In this way, the 8-oxo group would rescue the reduction in the T17 Hoogsteen exchange rate in hp^G•A^.

Indeed, for hp^8OG•A^ at T = 15°C, we did observe partial rescue of the T17 exchange kinetics (*k*ex ∼ 4200 s^-1^) which more closely mirrored values measured in the Watson-Crick hp^A-T^ and was roughly two-fold faster relative to hp^G•A^ (Fig. 3A-B). The modification had a smaller effect on the population (pop. ∼ 0.6%) and chemical shift difference (Δω ∼ −0.9 ppm) (Fig. 3B). Thus, the Watson-Crick-like 8OG*syn*-A*anti* conformation stabilized by the 8-oxoguanine lesion rescued the A-T breathing dynamics at a site that was two bps away. Once again, we did not observe any sign of exchange for the G-C bps.

Comparison of 2D HSQC spectra of the hairpin constructs also revealed that G•A in hp^G•A^ induced significant chemical shift perturbations (CSP) up to two bps away relative to the Watson-Crick hp^A-T^ for the aromatic resonances belonging to A3, A5, C6, G13, C16, T17 and for the sugar resonances of A5, C16, and T17. The 8OG•A in hp^8OG•A^ partially rescued these CSPs resulting in chemical shifts more like those of the Watson-Crick hp^A-T^ control (Fig. 3C and S9). Therefore, by impacting the 3D structure, mismatches, and lesions can affect the dynamics of distant bps. Together, these results demonstrate the power of ^1^H CEST to assess the impact of lesions comprehensively and systematically with respect to DNA dynamics not only on the affected bp but also on neighboring bps.

## Conclusion

This work establishes the utility of high-power ^1^H CEST as a fast and economic means for measuring conformational exchange in damaged nucleotides, which can be difficult to isotopically enrich and to also assess how the damage shapes the dynamics of neighboring bps in the duplex. Our findings lend further support that 8OG•A transiently forms a non-mutagenic 8OG*anti*•A*anti* conformation by enabling the direct measurement of the 8OG base flipping from the *syn* to the *anti* conformation. Our results also demonstrate that while the pyrimidine-pyrimidine mismatches that constrict C1ʹ-C1ʹ distance promote Watson-Crick to Hoogsteen breathing, purine-purine mismatches with wider C1ʹ-C1ʹ distances such as G•A slow down these same motions. Mismatches can also impact the dynamics of distant bps and be modulated by lesions such as 8OG, which also alter the C1ʹ-C1ʹ distance. We anticipate that the ^1^H CEST experiment can be used to comprehensively measure the sequence-dependence of 8OG•A dynamics as well as the dynamics of other damaged and/or modified DNA bases with unique proton signatures, including for 8OG- H1 when partnered with cytosine, pyrimidine dimers, those containing 5-methylcytosine, N6- methyladenosine, as well as DNA nicks.

## ASSOCIATED CONTENT

The following files are available free of charge.

Simulated ^1^H CEST, collected ^1^H CEST profiles, degeneracy analysis, and model selection analysis of ^1^H CEST, NMR spectra, collected data summary (PDF)

## Funding Sources

This work was supported by the NIH National Institute of General Medical Sciences (R01GM089846). The data collected at NYSBC using the 700 MHz spectrometer and 900 MHz NEO spectrometers was supported by the ORIP/NIH (CO6RR015495), the NIH (S10OD018509, P41GM066354, and S10OD030373), and the New York State Assembly. Some of the work presented here was conducted at the Center on Macromolecular Dynamics by NMR Spectroscopy located at NYSBC, supported by a grant from the NIH National Institute of General Medical Sciences RM1GM145397.

## Notes

The authors declare no competing financial interest.

## ACKNOWLEDGMENT

We would like to thank Ainan Geng (Duke University), Or Szekely (Duke University) and members of the Al-Hashimi lab for their input. We acknowledge resources from the Duke Magnetic Resonance Spectroscopy Center and the Duke Compute Cluster. H.M.A is a member of the New York Structural Biology Center (NYSBC).

## ABBREVIATIONS

8OG: 8-oxoguanine
AIC: Akaike’s information criterion
bp: base pair
BIC: Bayesian information criterion
B-M: Bloch-McConnell
CEST: Chemical exchange saturation transfer
CSP: Chemical shift perturbation
ES: excited state
GS: ground state
HSQC: Heteronuclear Single Quantum Coherence
NMR: Nuclear magnetic resonance
NOE: Nuclear Overhauser Effect
NOESY: Nuclear Overhauser Effect Spectroscopy
pop: population
*R*_1π_: spin relaxation in the rotating frame

## For Table of Contents Only

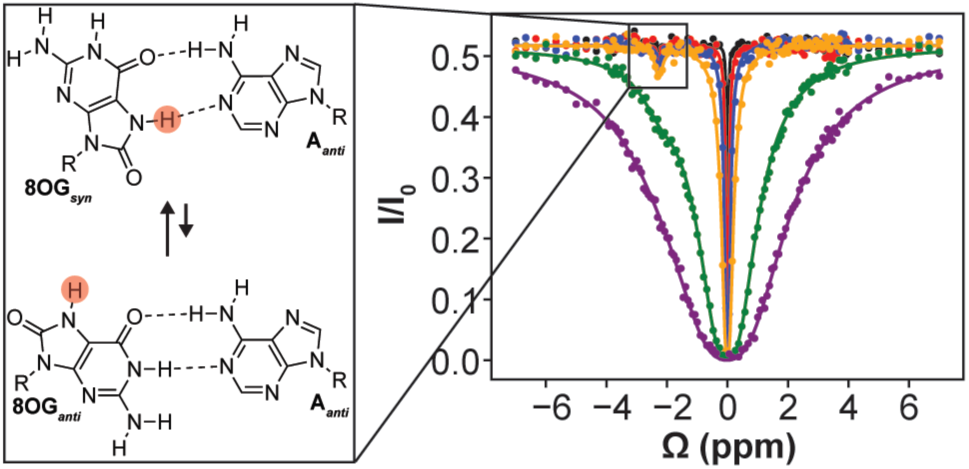

## Notes

### Competing Interest Statement

The authors have declared no competing interest.

