## Supplementary Info for "Direct Measurement of 8OG *syn-anti* Flips in Mutagenic 8OG•A and Long-Range Damage-Dependent Hoogsteen Breathing Dynamics Using ^1^H CEST NMR"

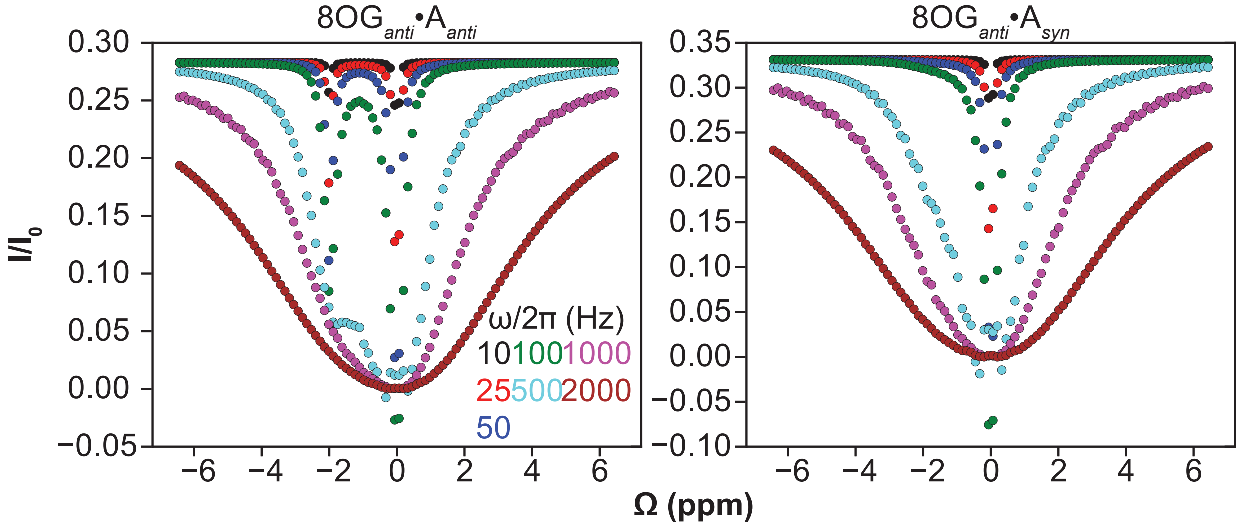

**Supplementary Figure 1. Simulated $^1$H CEST profiles for exchange involving 8OG$_{anti}$•A$_{anti}$ and 8OG$_{anti}$•A$_{syn}$ based on exchange parameters reported previously[1] using $^{13}$C $R_{1\rho}$ and CEST.** Simulated 8OG-H7 $^1$H CEST profiles assuming 2-state exchange between the 8OG$_{syn}$•A$_{anti}$ ground state and either 8OG$_{anti}$•A$_{anti}$ (assuming pop. = 15%; $k_{ex}$ = 200 s$^{-1}$; Δω = -2 ppm) or 8OG$_{anti}$•A$_{syn}$ (assuming pop. = 0.5%; $k_{ex}$ = 4700 s$^{-1}$; Δω = -2 ppm) as a minor excited state conformation. The exchange parameters were obtained from Gu *et al*[1] and were determined using $^{13}$C $R_{1\rho}$ and CEST experiments. Simulations show the much weaker contribution from the fast exchanging and lowly populated 8OG$_{anti}$•A$_{syn}$.

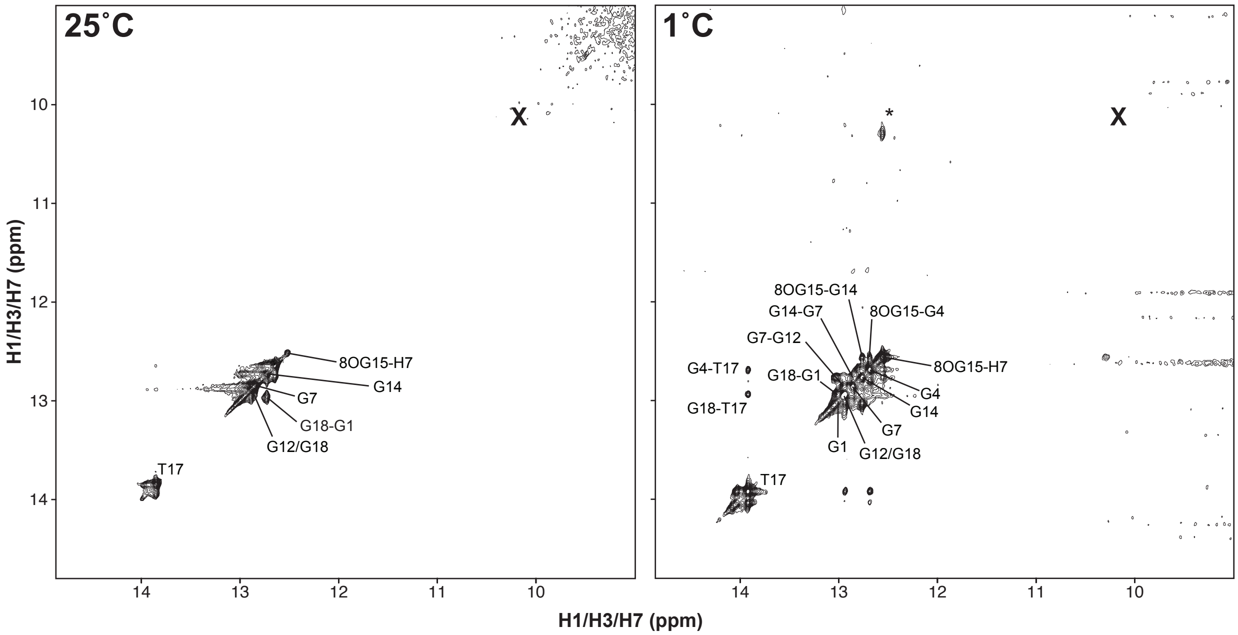

**Supplementary Figure 2. Temperature-dependent 2D [$^1$H-$^1$H] NOESY assignment of the imino region of hp$^{8OG•A}$.** No cross-peaks are observed within 4 ppm of the 8OG-H7 resonance. While a cross-peak was observed at ~10.3 ppm at 1˚C (marked with *), this did not correspond to any dips in the $^1$H CEST profiles at 10-25˚C. This cross-peak may be the NOE between 8OG-H7 and A-H6. The X denotes the expected diagonal peak position of the 8OG-H7 excited state conformation.

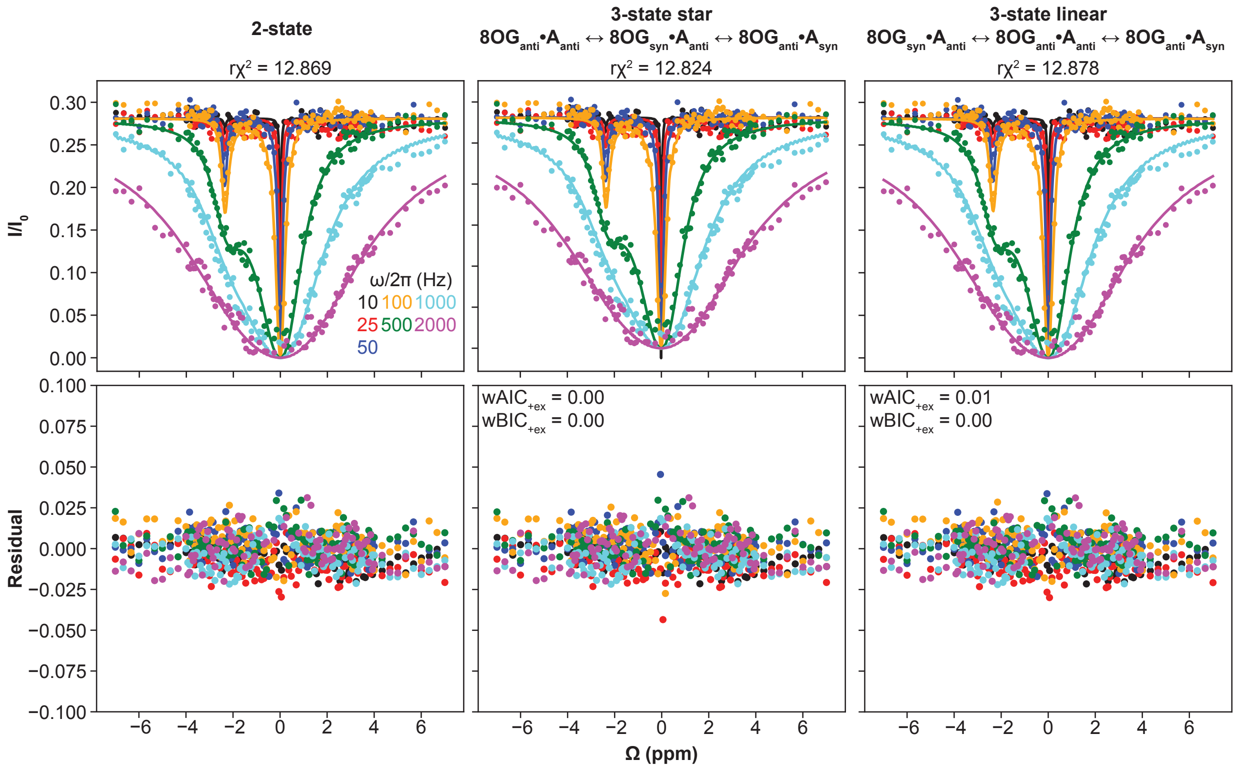

**Supplementary Figure 3. AIC/BIC model selection favors 2-state exchange for $8OG_{anti}$•$A_{anti}$.** AIC/BIC model selection[2-3] comparing 3-state star-like exchange and 3-state linear exchange to 2-state exchange favors the 2-state exchange. Data were collected at pH 7.4 and T = 25˚C.

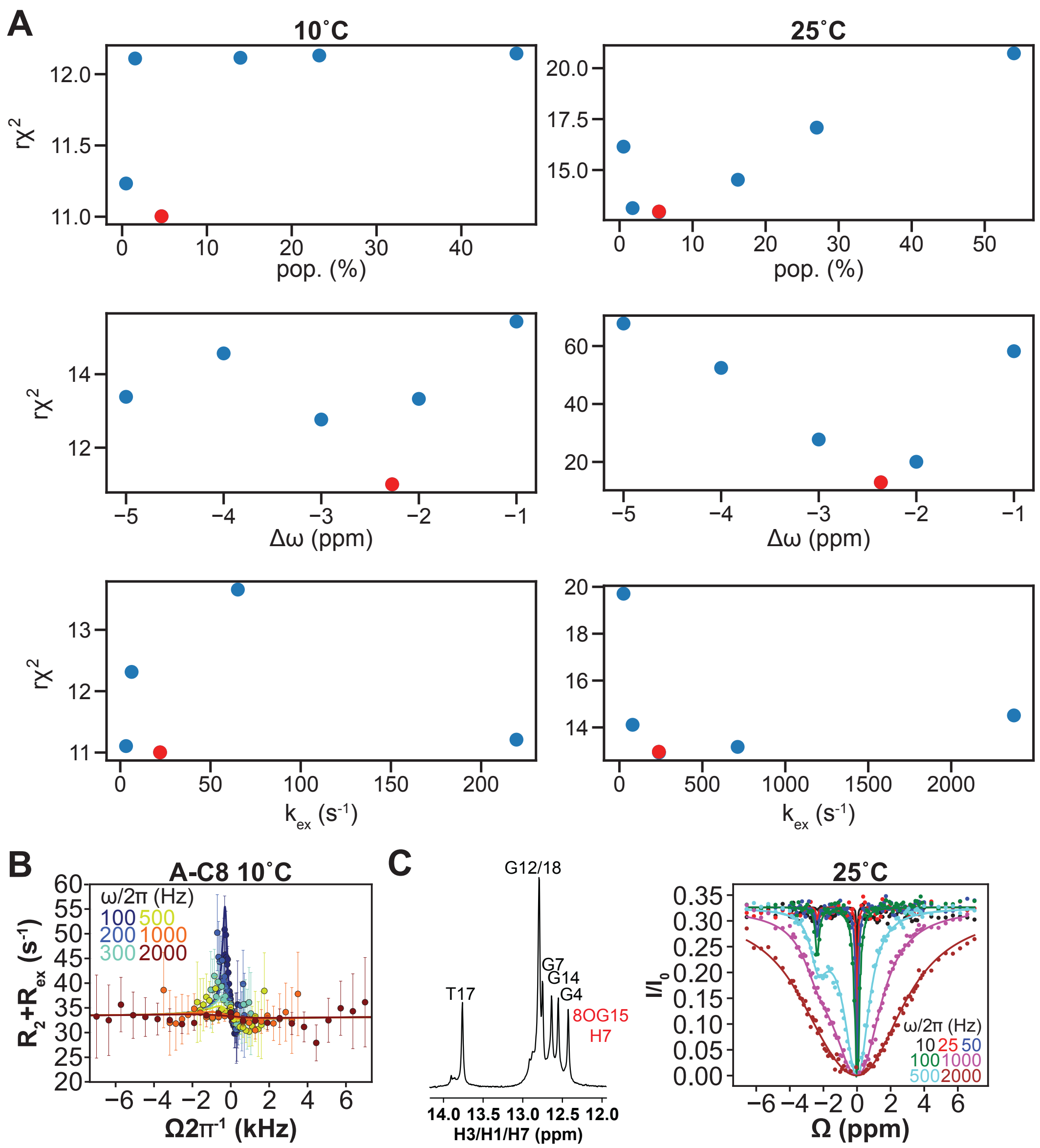

**Supplementary Figure 4. Additional data evaluating exchange involving 8OG$_{anti}$•A$_{anti}$ as an ES.** (A) Degeneracy analysis of $^{1}$H CEST data measured at 10˚C and 25˚C. Shown is the quality of a 2-state fit (r$\chi^2$) to the $^{1}$H CEST profile measured for 8OG-H7 when individually fixing each exchange parameter (pop., $k_{ex}$, or Δω) to a different value up to 10-fold or 3 ppm from the best-fit value while allowing all other exchange parameters to float during the fit. The best-fit exchange parameters are in red. (B) Off-resonance $^{13}$C $R_{1\rho}$ measurements on the adenine-C8 of a labeled hp$^{8OG•A}$ at T = 10˚C and same buffer conditions. Spin-lock powers are color coded. Error bars in the profile were obtained using Monte-Carlo simulations. The solid lines denote the fit of the data points to the B-M equations[4] assuming a 2-state exchange model. (C) Shown is the 1D imino region of hp$^{8OG•A}$ at T = 25˚C. The 8OG-H7 imino resonance is highlighted in red. Also shown is the 8OG-H7 $^{1}$H CEST profile measured at a field strength of 900 MHz at T = 25˚C as well as the fits to the $^{1}$H CEST data to two-state exchange using B-M equations with 2-state exchange. The error bars for the $^{1}$H CEST profiles, which are smaller than the data points, were obtained using triplicate experiments. RF powers are color-coded.

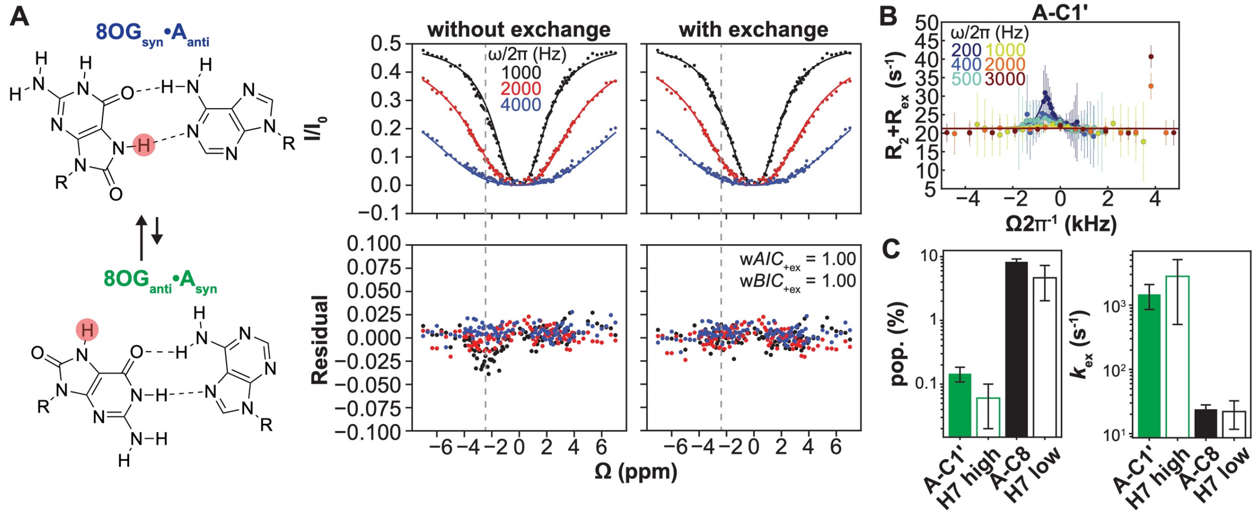

**Supplementary Figure 5. Detection of exchange consistent with 8OG$_{anti}$•A$_{syn}$ in hp$^{8OG•A}$ using $^{1}$H CEST at T = 10˚C**. (A) $^{1}$H CEST profiles measured using high spin-lock powers from 1,000 Hz to 4,000 Hz. The dashed gray lines indicate the ES $\Delta\omega$ position. The error bars for the $^{1}$H CEST profiles, which are smaller than the data points, were obtained using triplicate experiments. RF powers are color-coded. Shown are the fits of the $^{1}$H CEST data to a 2-state fit using B-M equations[4] with and without ($k_{ex}$ = $\Delta\omega$ = pop. = 0) chemical exchange. Shown below the profiles are residual plots (experimental normalized intensity – fitted normalized intensity). Also shown in the inset are Akaike's (w*AIC*) and Bayesian information criterion (w*BIC*) weights for fits with exchange. (B) $^{13}$C $R_{1\rho}$ measurements on the adenine-C1′ which is sensitive to the 8OG$_{anti}$•A$_{syn}$ conformational state measured in the labeled hp$^{8OG•A}$ at the same temperature and buffer conditions. RF powers are color-coded. The error bars for the $^{13}$C $R_{1\rho}$ profile were obtained from Monte Carlo simulations for one measurement. The solid lines denote the fit of the data points to the B-M equations assuming a 2-state exchange model. (C) Comparison of the best fit parameters between $^{1}$H CEST and $^{13}$C $R_{1\rho}$ indicate that the detected ES from $^{1}$H CEST is most likely 8OG$_{anti}$•A$_{syn}$ (green), and distinct from 8OG$_{anti}$•A$_{anti}$ (black) with respect to both pop. and $k_{ex}$ and in good agreement with exchange parameters determined via $^{13}$C $R_{1\rho}$ measurements. Errors correspond to fitting errors of $^{1}$H CEST, which were calculated as the square root of the diagonal elements of the covariance matrix, and $^{13}$C $R_{1\rho}$, which were calculated using a Monte-Carlo scheme.

### hp$^{A\text{-}T}$ 25˚C

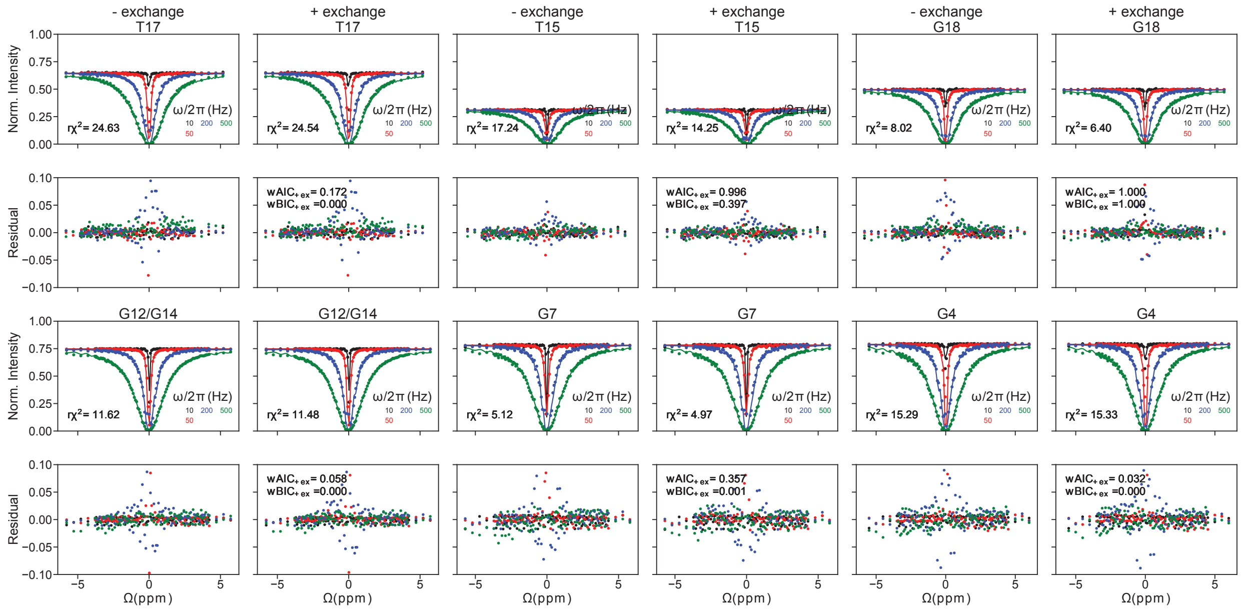

## hp$^{8OG•A}$ 25˚C

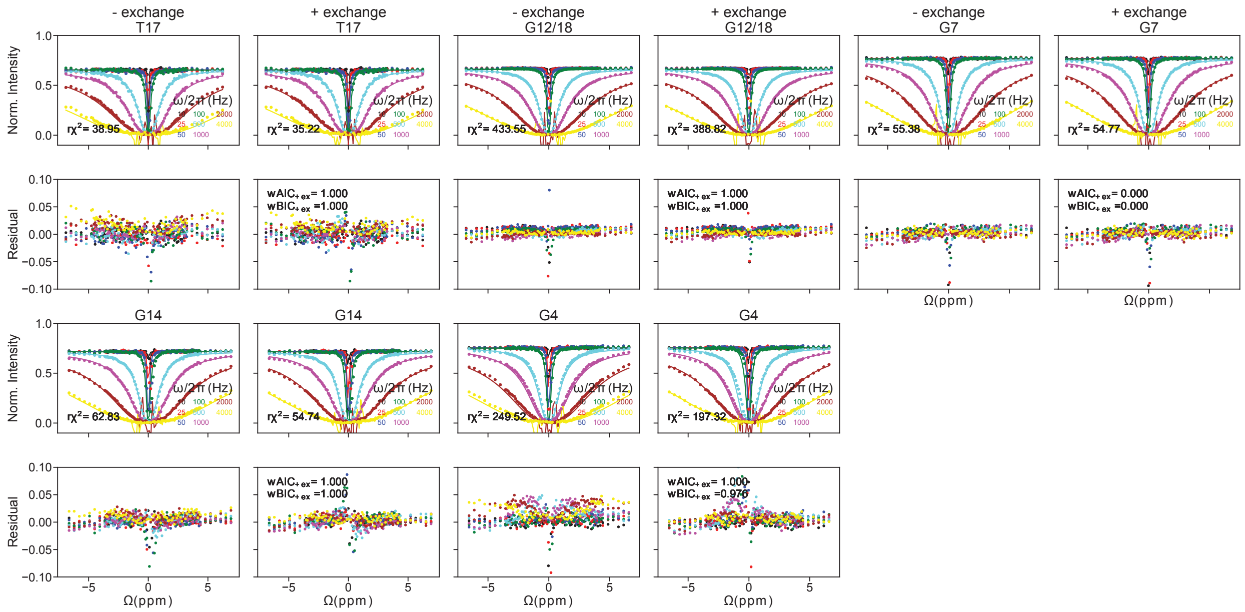

hp$^{A-T}$ 15˚C

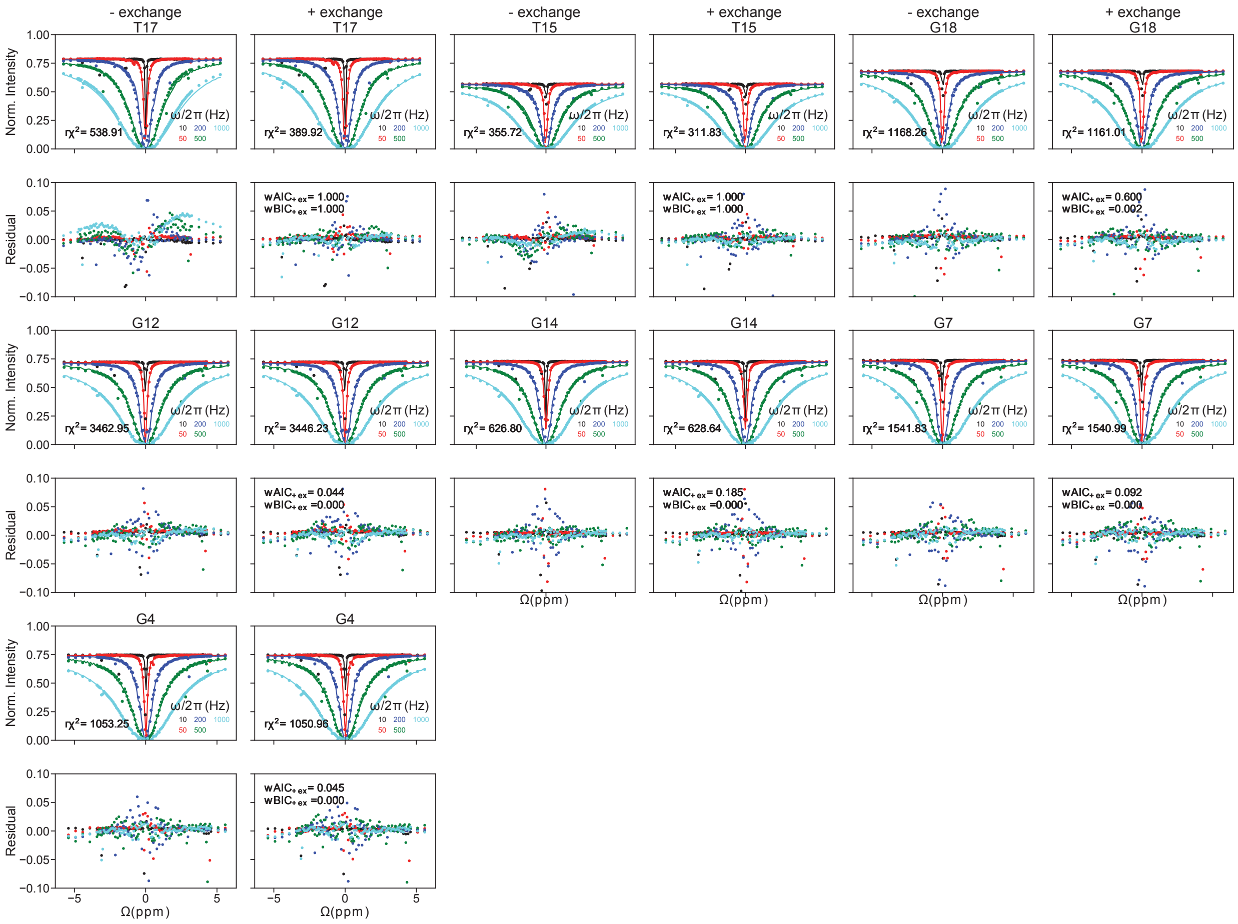

hp$^{8OG•A}$ 15˚C

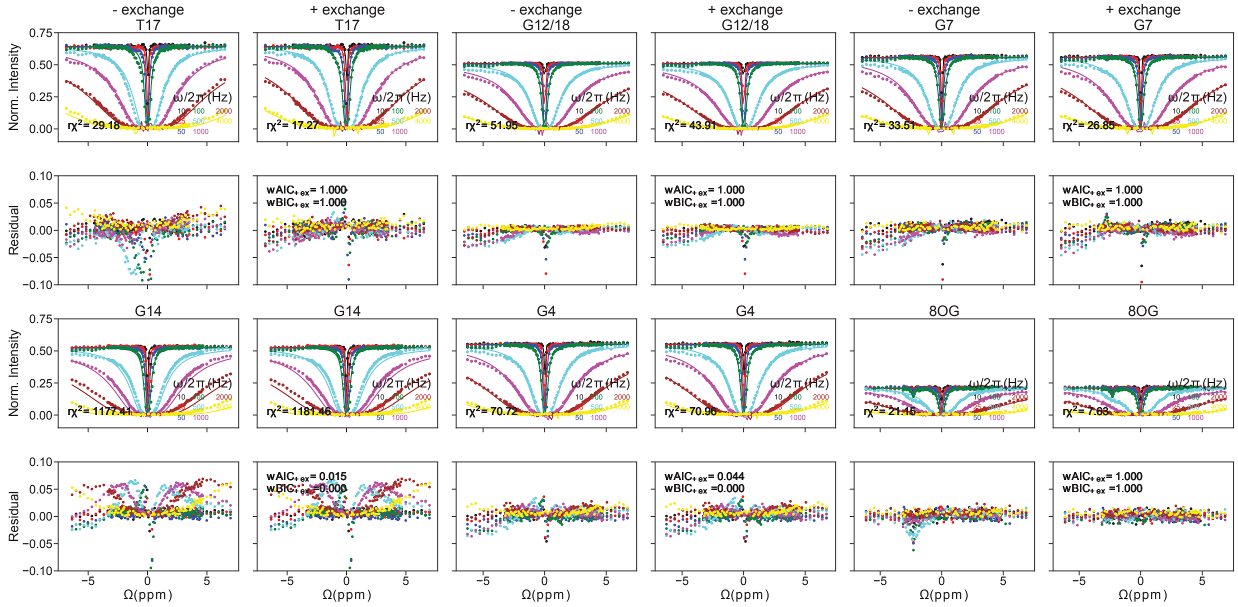

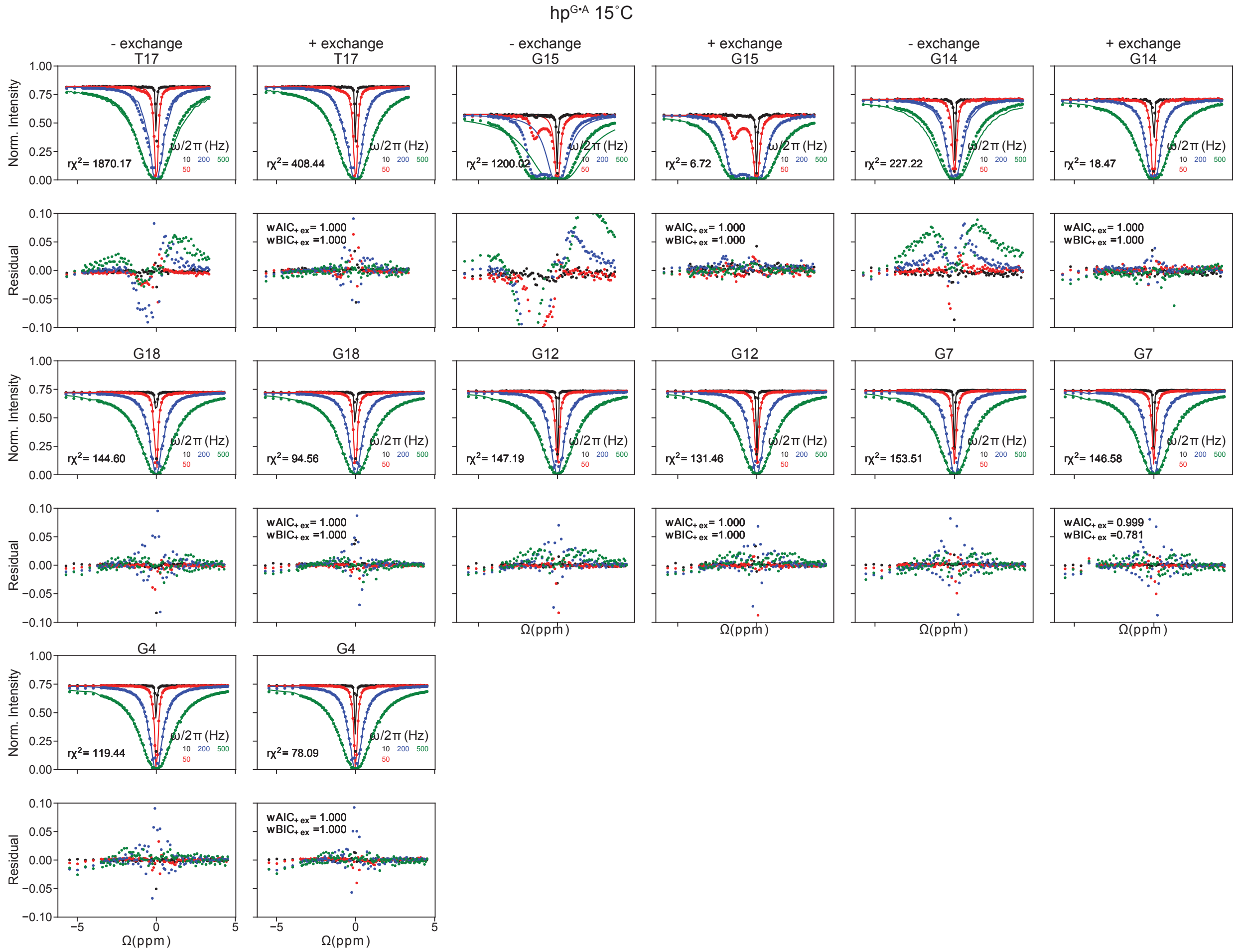

**Supplementary Figure 6. ¹H CEST profiles measured for imino protons at 25˚C and 15˚C.** Shown are the fits to the ¹H CEST data using B-M equations with and without 2-state chemical exchange. Below the ¹H CEST profiles are the residual plots. Also shown in inset are $r\chi^2$ and AIC/BIC weights for fits with exchange. RF powers are color-coded. Errors correspond to fitting errors of ¹H CEST, which were calculated as the square root of the diagonal elements of the covariance matrix.

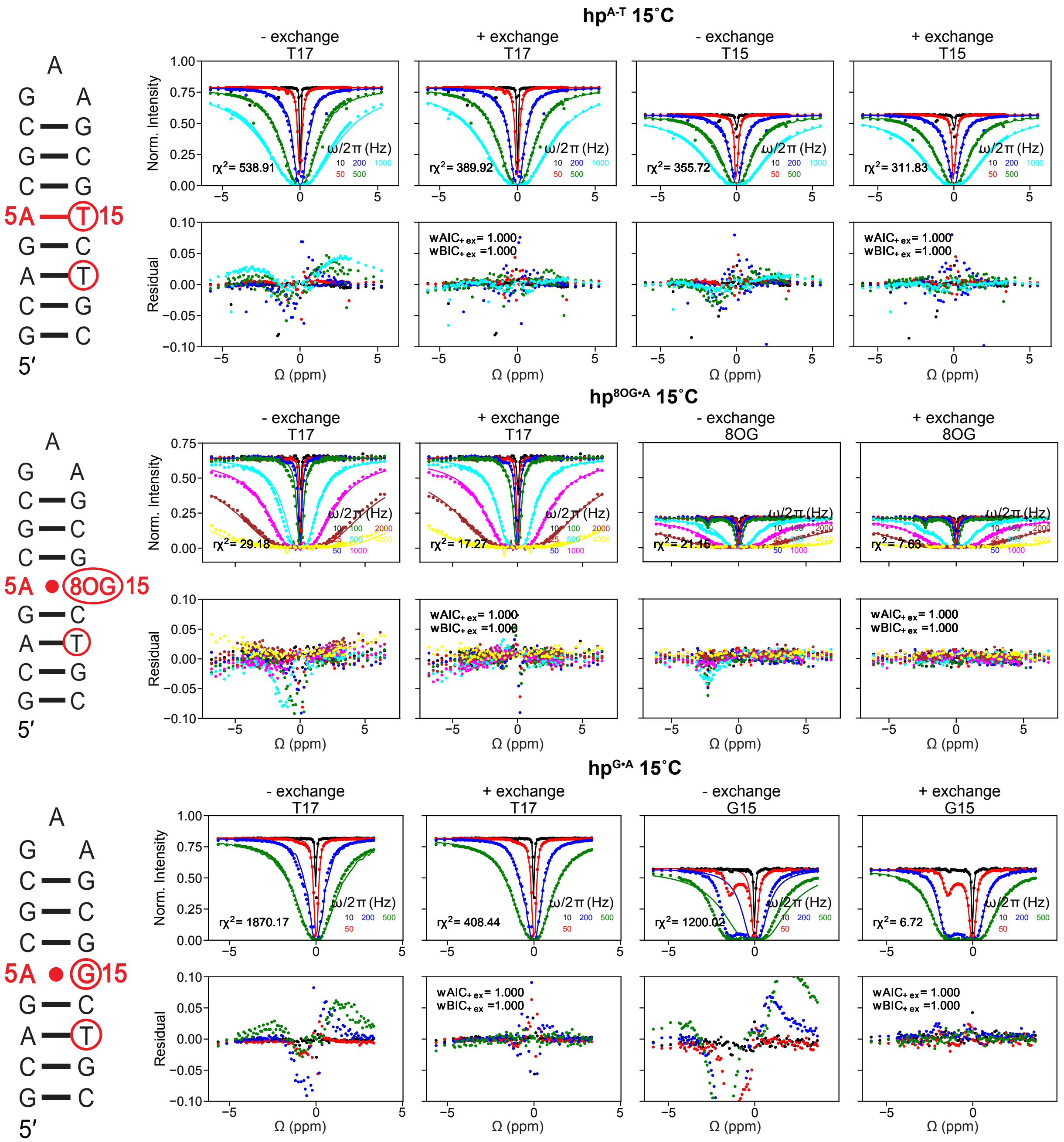

**Supplementary Figure 7. $^1$H CEST profiles measured for imino protons with an exchange contribution in hp$^{A\text{-}T}$, hp$^{A\bullet G}$, and hp$^{A\bullet 8OG}$ at T = 15˚C.** Resonances with detectable exchange are highlighted in red circles. Shown are the fits to the $^1$H CEST data using B-M equations with and without 2-state chemical exchange. Shown below the $^1$H CEST profiles are the residual plots (experimental normalized intensity – fitted normalized intensity). Also shown in inset are r$\chi^2$ and Akaike's (w*AIC*) and Bayesian information criterion (w*BIC*) weights for fits with exchange. The error bars for the $^1$H CEST profiles, which are smaller than the data points, were obtained using triplicate experiments. RF powers are color-coded. Errors correspond to fitting errors of $^1$H CEST, which were calculated as the square root of the diagonal elements of the covariance matrix.

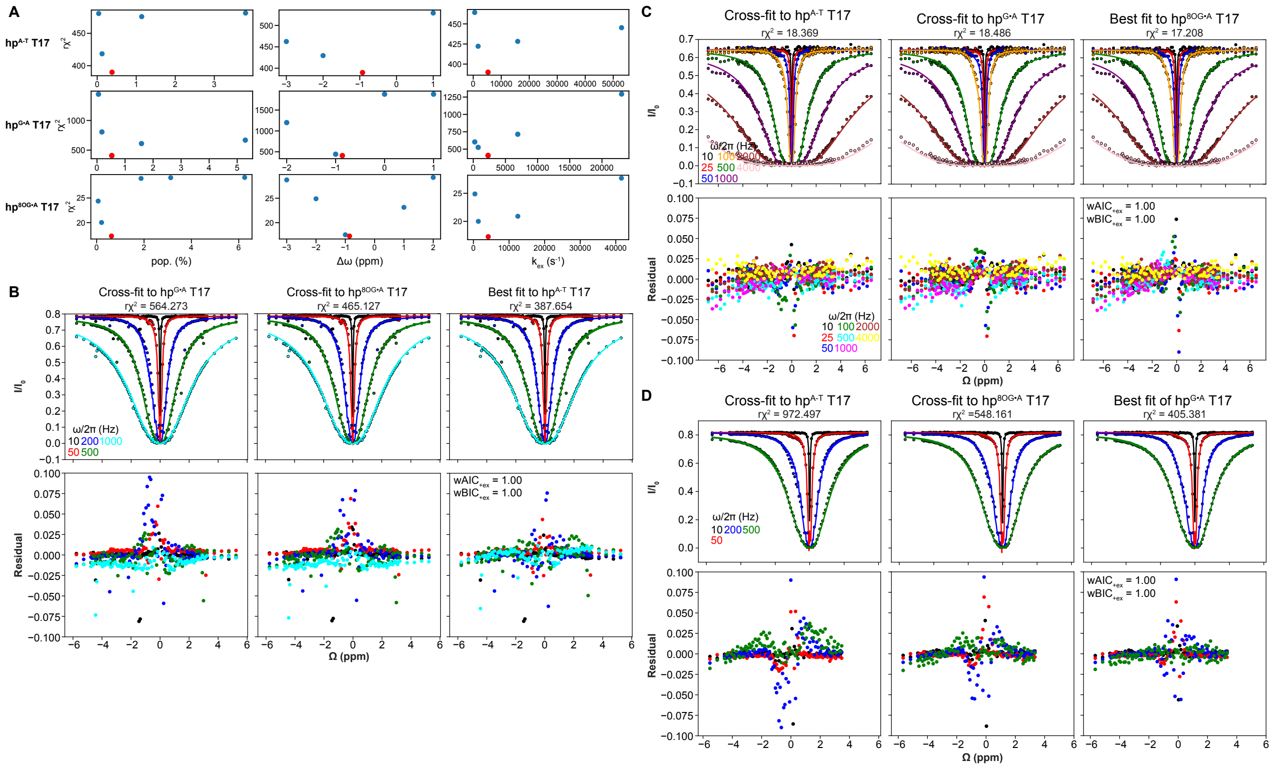

**Supplementary Figure 8. Testing statistical significance of exchange parameters deduced using $^1$H CEST for T17 at 15˚C.** (A) The quality of a 2-state fit ($r\chi^2$) to the $^1$H CEST profile measured for T17-H3 as a function of individually fixing the exchange parameters (pop., $k_{ex}$, or Δω) to a different value while allowing all other exchange parameters to float during the fit of the $^1$H CEST profile. The best-fit exchange parameters are in red. (B) $^1$H CEST profiles of $hp^{A\text{-}T}$ T17 data fixed to exchange parameters (pop., $k_{ex}$, Δω) corresponding to those of $hp^{G•A}$, $hp^{8OG•A}$, and its best fit. Shown below the $^1$H CEST profiles are the residual plots. Also shown in inset are $r\chi^2$, AIC, and BIC weights for fits compared to the $hp^{A\text{-}T}$ T17 best fit exchange parameters. (C) $^1$H CEST profiles of $hp^{8OG•A}$ T17 data fixed to exchange parameters corresponding to those of $hp^{A\text{-}T}$, $hp^{G•A}$, and its best fit. Shown below the $^1$H CEST profiles are the residual plots. Also shown in inset are $r\chi^2$, AIC, and BIC weights for fits compared to the $hp^{8OG•A}$ T17 best fit exchange parameters. (D) $^1$H CEST profiles of $hp^{G•A}$ T17 data fixed to exchange parameters corresponding to those of $hp^{A\text{-}T}$, $hp^{8OG•A}$, and its best fit. Shown below the $^1$H CEST profiles are the residual plots. Also shown in inset are $r\chi^2$, AIC, and BIC weights for fits compared to the $hp^{A\text{-}T}$ T17 best fit exchange parameters. The error bars for the $^1$H CEST profiles, which are smaller than the data points, were obtained using triplicate experiments. RF powers are color-coded.

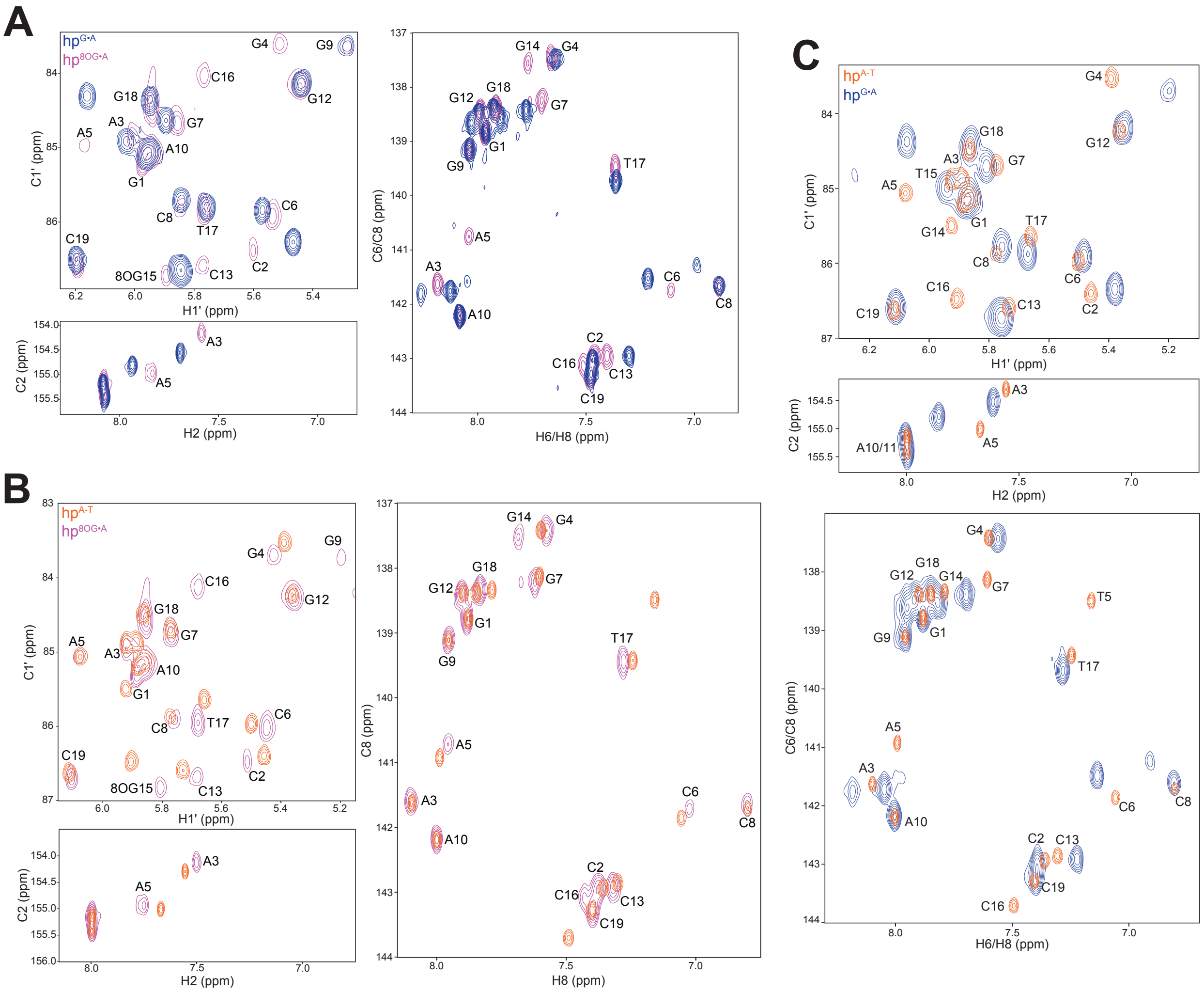

**Supplementary Figure 9. Overlay of aromatic C2H2/C6H6/C8H8 and sugar C1′H1′ 2D HSQC spectra measured for the three hairpins in this study at T = 25˚C.** (A) Comparison between hp$^{G\bullet A}$ (blue) and hp$^{8OG\bullet A}$ (purple) with the assignments for hp$^{8OG\bullet A}$ shown. (B) Comparison between hp$^{A\text{-}T}$ (red) and hp$^{8OG\bullet A}$ (purple) with the assignments for hp$^{8OG\bullet A}$ shown. (C) Comparison between hp$^{A\text{-}T}$ (red) and hp$^{G\bullet A}$ (blue) with peak assignments for hp$^{A\text{-}T}$ shown.

**Supplementary Table 1.** List of spin-lock powers ($\omega_1/2\pi$, in Hz) and offsets ($\Omega/2\pi$, in Hz) used in $^{1}$H CEST experiments and RD experiments.

| Nucleus | [$\omega_1/2\pi$ (Hz)] [$\Omega/2\pi$ (Hz)] |
|---|---|
| hp$^{8OG\bullet A}$ (pH 7.4, 25°C, 90% $H_2O$:10% $D_2O$) on Bruker 700 MHz | |
| 8OG15 H7 | [10, 25, 50, 100, 500, 1000, 2000] [-4896.827426, -4663.404326, -4429.980526, -4196.557426, -3963.134326, -3729.710526, -3496.287426, -3262.864326, -3029.440526, -2796.017426, -2742.150557, -2688.283688, -2634.416819, -2580.549949, -2526.68308, -2472.815511, -2418.949342, -2365.081772, -2311.214903, -2257.348034, -2203.481165, -2149.614296, -2095.747426, -2041.880557, -1988.013688, -1934.146819, -1880.279949, -1826.41308, -1772.545511, -1718.679342, -1664.811772, -1610.944903, -1557.078034, -1503.211165, -1449.344296, -1395.477426, -1341.610557, -1287.743688, -1233.876819, -1180.009949, -1126.14308, -1072.276211, -1018.408642, -964.5417724, -910.6749032, -856.808034, -802.9411648, -749.0742955, -695.2074263, -621.4947653, -547.7821742, -474.0695132, -400.3569221, -326.6442611, -252.9316, -179.219009, -105.5063479, -31.79374286, 41.91889019, 115.6314952, 189.3441563, 263.0567473, 336.7694084, 410.4820695, 484.1946605, 557.9073216, 631.6199126, 705.3325737, 759.1994429, 813.0663121, 866.9331813, 920.8000505, 974.6669197, 1028.533789, 1082.401358, 1136.268228, 1190.135097, 1244.001966, 1297.868835, 1351.735704, 1405.602574, 1459.469443, 1513.336312, 1567.203181, 1621.070051, 1674.93692, 1728.804489, 1782.670658, 1836.538228, 1890.405097, 1944.271966, 1998.138835, 2052.005704, 2105.872574, 2159.739443, 2213.606312, 2267.473181, 2321.340051, 2375.20692, 2429.074489, 2482.940658, 2536.808228, 2590.675097, 2644.541966, 2698.408835, 2752.275704, 2806.142574, 3039.565674, 3272.989474, 3506.412574, 3739.835674, 3973.259474, 4206.682574, 4440.105674, 4673.529474, 4906.952574] |
| hp$^{8OG\bullet A}$ (pH 7.4, 15˚C, 90% $H_2O$:10% $D_2O$) on Bruker 900 MHz | |
| All iminos | [10, 25, 50, 100, 500, 1000, 2000, 4000] [-6300.679685, -6000.603318, -5700.526051, -5400.449685, -5100.373318, -4800.296051, -4500.219685, -4200.143318, -3900.066051, -3599.989685, -3530.741292, -3461.4929, -3392.244508, -3322.996116, -3253.747723, -3184.498431, -3115.250939, -3046.001646, -2976.753254, -2907.504862, -2838.256469, -2769.008077, -2699.759685, -2630.511292, -2561.2629, -2492.014508, -2422.766116, -2353.517723, -2284.268431, -2215.020939, -2145.771646, -2076.523254, -2007.274862, -1938.026469, -1868.778077, -1799.529685, -1730.281292, -1661.0329, -1591.784508, -1522.536116, -1453.287723, -1384.039331, -1314.790038, -1245.541646, -1176.293254, -1107.044862, -1037.796469, -968.548077, -899.2996847, -804.5385941, -709.7775936, -615.0165031, -520.2555026, -425.494412, -330.7333215, -235.972321, -141.2112304, -46.45021193, 48.3108426, 143.0718611, 237.8329517, 332.5939522, 427.3550427, 522.1161332, 616.8771337, 711.6382243, 806.3992248, 901.1603153, 970.4087076, 1039.6571, 1108.905492, 1178.153884, 1247.402277, |

| | |
|---|---|
| | 1316.650669, 1385.899962, 1455.148354, 1524.396746, 1593.645138, 1662.893531, 1732.141923, 1801.390315, 1870.638708, 1939.8871, 2009.135492, 2078.383884, 2147.632277, 2216.881569, 2286.129061, 2355.378354, 2424.626746, 2493.875138, 2563.123531, 2632.371923, 2701.620315, 2770.868708, 2840.1171, 2909.365492, 2978.613884, 3047.862277, 3117.111569, 3186.359061, 3255.608354, 3324.856746, 3394.105138, 3463.353531, 3532.601923, 3601.850315, 3901.926682, 4202.003949, 4502.080315, 4802.156682, 5102.233949, 5402.310315, 5702.386682, 6002.463949, 6302.540315] |
| hp$^{8OG\bullet A}$ (pH 7.4, 10˚C, 90% $H_2O$:10% $D_2O$) on Bruker 900 MHz | |
| 8OG15 H7 | [10, 25, 50, 100, 500, 1000, 2000, 4000] [-6301.041353 , -6000.964987, -5700.88772, -5400.811353, -5100.734987, -4800.65772, -4500.581353, -4200.504987, -3900.42772, -3600.351353, -3531.102961, -3461.854569, -3392.606176, -3323.357784, -3254.109392, -3184.860099, -3115.612607, -3046.363315, -2977.114922, -2907.86653, -2838.618138, -2769.369746, -2700.121353, -2630.872961, -2561.624569, -2492.376176, -2423.127784, -2353.879392, -2284.630099, -2215.382607, -2146.133315, -2076.884922, -2007.63653, -1938.388138, -1869.139746, -1799.891353, -1730.642961, -1661.394569, -1592.146176, -1522.897784, -1453.649392, -1384.401, -1315.151707, -1245.903315, -1176.654922, -1107.40653, -1038.158138, -968.9097456, -899.6613533, -804.9002628, -710.1392623, -615.3781717, -520.6171712, -425.8560807, -331.0949902, -236.3339897, -141.5728991, -46.8118806, 47.94917393, 142.7101924, 237.471283, 332.2322835, 426.993374, 521.7544646, 616.5154651, 711.2765556, 806.0375561, 900.7986467, 970.047039, 1039.295431, 1108.543824, 1177.792216, 1247.040608, 1316.289, 1385.538293, 1454.786685, 1524.035078, 1593.2834, 1662.531862, 1731.780254, 1801.028647, 1870.277039, 1939.525431, 2008.773824, 2078.022216, 2147.270608, 2216.519901, 2285.767393, 2355.016685, 2424.265078, 2493.51347, 2562.761862, 2632.010254, 2701.258647, 2770.507039, 2839.755431, 2909.003824, 2978.252216, 3047.500608, 3116.749901, 3185.997393, 3255.246685, 3324.495078, 3393.74347, 3462.991862, 3532.240254, 3601.488647, 3901.565013, 4201.64228, 4501.718647, 4801.795013, 5101.87228, 5401.948647, 5702.025013, 6002.10228, 6302.178647] |
| hp$^{8OG\bullet A}$ (pH 7.4, 25˚C, 90% $H_2O$:10% $D_2O$) on Bruker 900 MHz | |
| 8OG15 H7 | [10, 25, 50, 100, 500, 1000, 2000] [-6303.394583, -5917.582112, -5531.76874, -5145.956269, -4760.142897, -4374.330426, -3988.517054, -3602.704583, -3509.57759, -3416.449697, -3323.322704, -3230.195711, -3137.068718, -3043.940824, -2950.813831, -2857.686838, -2764.558945, -2671.431952, -2578.304959, -2485.177966, -2392.050073, -2298.92308, -2205.796086, -2112.669093, -2019.5412, -1926.414207, -1833.287214, -1740.159321, -1647.032328, -1553.905335, -1460.778342, -1367.650448, -1274.523455, -1181.396462, -1088.269469, -995.141576, -902.014583, -807.2534925, -712.492492, -617.7314014, -522.9704009, -428.2093104, -333.4482198, -238.6872193, -143.9261288, -49.16511026, 45.59594426, 140.3569628, 235.1180533, 329.8790538, 424.6401444, 519.4012349, 614.1622354, |

| | |
|---|---|
| | 708.923326, 803.6843265, 898.445417, 991.57241, 1084.700303, 1177.827296, 1270.954289, 1364.081282, 1457.209176, 1550.336169, 1643.463162, 1736.590155, 1829.718048, 1922.845041, 2015.972034, 2109.099927, 2202.22692, 2295.353914, 2388.480907, 2481.6088, 2574.735793, 2667.862786, 2760.989779, 2854.117672, 2947.244665, 3040.371658, 3133.499552, 3226.626545, 3319.753538, 3412.880531, 3506.008424, 3599.135417, 3984.947888, 4370.76126, 4756.573731, 5142.387103, 5528.199574, 5914.012946, 6299.825417] |
| hp$^{8OG•A}$ (pH 7.4, 25˚C, 90% $H_2O$:10% $D_2O$) on Bruker 900 MHz | |
| G-H1 and T-H3 | [10, 25, 50, 100, 500, 1000, 2000, 4000] [-6990.981, -6605.16852867, -6219.35515711, -5833.54268578, -5447.72931422, -5061.91684289, -4676.10347133, -4290.291, -4197.16400696, -4104.03611369, -4010.90912065, -3917.78212761, -3824.65513457, -3731.5272413, -3638.40024826, -3545.27325522, -3452.14536195, -3359.01836891, -3265.89137587, -3172.76438283, -3079.63648956, -2986.50949652, -2893.38250348, -2800.25551044, -2707.12761717, -2614.00062413, -2520.87363109, -2427.74573782, -2334.61874478, -2241.49175174, -2148.3647587, -2055.23686543, -1962.10987239, -1868.98287935, -1775.85588631, -1682.72799304, -1589.601, -1494.83990946, -1400.07890895, -1305.31781842, -1210.5568179, -1115.79572737, -1021.03463683, -926.273636317, -831.512545781, -736.751527263, -641.990472737, -547.229454219, -452.468363683, -357.70736317, -262.946272634, -168.185182098, -73.424181585, 116.097909464, 210.859, 303.98599304, 397.11388631, 490.24087935, 583.36787239, 676.49486543, 769.6227587, 862.74975174, 955.87674478, 1049.00373782, 1142.13163109, 1235.25862413, 1328.38561717, 1421.51351044, 1514.64050348, 1607.76749652, 1700.89448956, 1794.02238283, 1887.14937587, 1980.27636891, 2073.40336195, 2166.53125522, 2259.65824826, 2352.7852413, 2445.91313457, 2539.04012761, 2632.16712065, 2725.29411369, 2818.42200696, 2911.549, 3297.36147133, 3683.17484289, 4068.98731422, 4454.80068578, 4840.61315711, 5226.42652867, 5612.239] |
| hp$^{A\text{-}T}$ (pH 7.4, 25˚C, 90% $H_2O$:10% $D_2O$) on Bruker 600 MHz | |
| All iminos | [10, 50, 200, 500, 1000] [-4092.199, -3792.369, -3492.539, -3192.709, -2892.879, -2844.42167472, -2795.96434944, -2747.50702416, -2699.04969888, -2650.5923736, -2602.13504832, -2553.67712338, -2505.2197981, -2456.76247282, -2408.30514754, -2359.84782226, -2311.39049698, -2262.9331717, -2214.47584642, -2166.01852114, -2117.56119586, -2069.10387058, -2020.6465453, -1972.18922002, -1923.73129508, -1875.2739698, -1826.81664452, -1778.35931924, -1729.90199396, -1681.44466868, -1632.9873434, -1584.53001812, -1536.07269284, -1487.61536756, -1439.15804228, -1390.70011734, -1342.24279206, -1293.78546678, -1245.3281415, -1196.87081622, -1148.41349094, -1099.95616566, -1051.49878041, -1003.04145513, -954.584069888, -906.126684642, -857.669299396, -809.21191415, -760.754528904, -712.297203624, -663.839818378, -615.382433132, -566.925047886, -518.467686626, -470.010313374, -421.552952114, -373.095566868, - |

| | |
|---|---|
| | 324.638181622, -276.180796376, -227.723471096, -179.26608585, -130.808700604, -82.351315358, -33.893930112, 14.563455134, 63.020780414, 111.47816566, 159.93549094, 208.39281622, 256.8501415, 305.30746678, 353.76479206, 402.22211734, 450.68004228, 499.13736756, 547.59469284, 596.05201812, 644.5093434, 692.96666868, 741.42399396, 789.88131924, 838.33864452, 886.7959698, 935.25329508, 983.71122002, 1032.1685453, 1080.62587058, 1129.08319586, 1177.54052114, 1225.99784642, 1274.4551717, 1322.91249698, 1371.36982226, 1419.82714754, 1468.28447282, 1516.7417981, 1565.19912338, 1613.65704832, 1662.1143736, 1710.57169888, 1759.02902416, 1807.48634944, 1855.94367472, 1904.401, 2204.231, 2504.061, 2803.891, 3103.721] |
| hp$^{A-T}$ (pH 7.4, 15˚C, 90% $H_2O$:10% $D_2O$) on Bruker 600 MHz | |
| All iminos | [10, 50, 200, 500, 1000] [-4056.02766667, -3756.19766667, -3456.36766667, -3156.53766667, -2856.70766667, -2808.25034139, -2759.79301611, -2711.33569083, -2662.87836555, -2614.42104027, -2565.96371499, -2517.50579005, -2469.04846477, -2420.59113949, -2372.13381421, -2323.67648893, -2275.21916365, -2226.76183837, -2178.30451309, -2129.84718781, -2081.38986253, -2032.93253725, -1984.47521197, -1936.01788669, -1887.55996175, -1839.10263647, -1790.64531119, -1742.18798591, -1693.73066063, -1645.27333535, -1596.81601007, -1548.35868479, -1499.90135951, -1451.44403423, -1402.98670895, -1354.52878401, -1306.07145873, -1257.61413345, -1209.15680817, -1160.69948289, -1112.24215761, -1063.78483233, -1015.32744708, -966.870121801, -918.412736555, -869.955351309, -821.497966063, -773.040580817, -724.583195571, -676.125870291, -627.668485045, -579.211099799, -530.753714553, -482.296353293, -433.83898004, -385.381618781, -336.924233535, -288.466848289, -240.009463043, -191.552137763, -143.094752517, -94.6373672707, -46.1799820247, 50.7347884673, 99.1921137473, 147.649498993, 196.106824273, 244.564149553, 293.021474833, 341.478800113, 389.936125393, 438.393450673, 486.851375613, 535.308700893, 583.766026173, 632.223351453, 680.680676733, 729.138002013, 777.595327293, 826.052652573, 874.509977853, 922.967303133, 971.424628413, 1019.88255335, 1068.33987863, 1116.79720391, 1165.25452919, 1213.71185447, 1262.16917975, 1310.62650503, 1359.08383031, 1407.54115559, 1455.99848087, 1504.45580615, 1552.91313143, 1601.37045671, 1649.82838165, 1698.28570693, 1746.74303221, 1795.20035749, 1843.65768277, 1892.11500805, 1940.57233333, 2240.40233333, 2540.23233333, 2840.06233333, 3139.89233333] |
| hp$^{G•A}$ (pH 7.4, 15˚C, 90% $H_2O$:10% $D_2O$) on Bruker 600 MHz | |
| | [10, 50, 200, 500] [-3995.245, -3695.415, -3395.585, -3095.755, -2795.925, -2747.46767472, -2699.01034944, -2650.55302416, -2602.09569888, -2553.6383736, -2505.18104832, -2456.72312338, -2408.2657981, -2359.80847282, -2311.35114754, -2262.89382226, -2214.43649698, -2165.9791717, -2117.52184642, -2069.06452114, -2020.60719586, -1972.14987058, -1923.6925453, -1875.23522002, -1826.77729508, - |

| | |
|---|---|
| | 1778.3199698, -1729.86264452, -1681.40531924, -1632.94799396, -1584.49066868, -1536.0333434, -1487.57601812, -1439.11869284, -1390.66136756, -1342.20404228, -1293.74611734, -1245.28879206, -1196.83146678, -1148.3741415, -1099.91681622, -1051.45949094, -1003.00216566, -954.544780414, -906.087455134, -857.630069888, -809.172684642, -760.715299396, -712.25791415, -663.800528904, -615.343203624, -566.885818378, -518.428433132, -469.971047886, -421.513686626, -373.056313374, -324.598952114, -276.141566868, -227.684181622, -179.226796376, -130.769471096, -82.31208585, 111.517455134, 159.974780414, 208.43216566, 256.88949094, 305.34681622, 353.8041415, 402.26146678, 450.71879206, 499.17611734, 547.63404228, 596.09136756, 644.54869284, 693.00601812, 741.4633434, 789.92066868, 838.37799396, 886.83531924, 935.29264452, 983.7499698, 1032.20729508, 1080.66522002, 1129.1225453, 1177.57987058, 1226.03719586, 1274.49452114, 1322.95184642, 1371.4091717, 1419.86649698, 1468.32382226, 1516.78114754, 1565.23847282, 1613.6957981, 1662.15312338, 1710.61104832, 1759.0683736, 1807.52569888, 1855.98302416, 1904.44034944, 1952.89767472, 2001.355] |
| hp$^{8OG•A}$ (pH 7.4, 10˚C, 90% $H_2O$:10% $D_2O$) on Bruker 700 MHz | |
| A-C8 | [100] [-10, -32, -64, -96, -128, -160, -192, -224, -256, -288, -320, -352, 10, 32, 64, 96, 128, 160, 192, 224, 256, 288, 320, 352]<br>[200] [-10, -64, -128, -192, -256, -320, -384, -448, -512, -576, -640, -704, 10, 64, 128, 192, 256, 320, 384, 448, 512, 576, 640, 704]<br>[300] [-10, -95, -190, -285, -380, -475, -570, -665, -760, -855, -950, -1045, 10, 95, 190, 285, 380, 475, 570, 665, 760, 855, 950, 1045]<br>[500] [-10, -159, -318, -477, -636, -795, -954, -1113, -1272, -1431, -1590, -1749, 10, 159, 318, 477, 636, 795, 954, 1113, 1272, 1431, 1590, 1749]<br>[1000] [-10, -318, -636, -954, -1272, -1590, -1908, -2226, -2544, -2862, -3180, -3498, 10, 318, 636, 954, 1272, 1590, 1908, 2226, 2544, 2862, 3180, 3498]<br>[2000] [-10, -636, -1272, -1908, -2544, -3180, -3816, -4452, -5088, -5724, -6360, -6996, 10, 636, 1272, 1908, 2544, 3180, 3816, 4452, 5088, 5724, 6360, 6996] |
| A-C1′ | [200] [-10, -64, -128, -192, -256, -320, -384, -448, -512, -576, -640, -704, 10, 64, 128, 192, 256, 320, 384, 448, 512, 576, 640, 704], [400] [-10, -127, -254, -381, -508, -635, -762, -889, -1016, -1143, -1270, -1397, 10, 127, 254, 381, 508, 635, 762, 889, 1016, 1143, 1270, 1397]<br>[500] [-10, -159, -318, -477, -636, -795, -954, -1113, -1272, -1431, -1590, -1749, 10, 159, 318, 477, 636, 795, 954, 1113, 1272, 1431, 1590, 1749]<br>[1000] [-10, -318, -636, -954, -1272, -1590, -1908, -2226, -2544, -2862, -3180, -3498, 10, 318, 636, 954, 1272, 1590, 1908, 2226, 2544, 2862, 3180, 3498]<br>[2000] [-10, -636, -1272, -1908, -2544, -3180, -3816, -4452, -5088, -5724, -6360, -6996, 10, 636, 1272, 1908, 2544, 3180, 3816, 4452, 5088, 5724, 6360, 6996]<br>[3000] [-10, -955, -1910, -2865, -3820, -4775, -5730, -6685, -7640, -8595, -9550, -10505, 10, 955, 1910, 2865, 3820, 4775, 5730, 6685, 7640, 8595, 9550, 10505] |

**Supplementary Table 2.** Summary of exchange parameters obtained from fitting 8OG$_{anti}$•A$_{anti}$ (ES1) to $^{1}$H CEST data measured on unlabeled hp$^{8OG•A}$ 8OG-H7 across two temperatures at pH 7.4 and in 90% $H_2O$:10% $D_2O$. Red. $\chi^2$ denotes the reduced $\chi^2$ obtained on fitting the $^{1}$H CEST data. At T = 10˚C, the best fit parameters for 8OG$_{anti}$•A$_{syn}$ (ES2) were also included based on the high spin-lock power data.

| T = 25˚C | | |
|---|---|---|
| Parameter | 8OG-H7 700 MHz | 8OG-H7 900 MHz |
| $pop_{ES1}$ (%) | 5.1 ± 2.7 | 2.8 ±1.5 |
| $k_{ex,ES1}$ ($s^{-1}$) | 255 ± 64 | 304 ± 49 |
| $\Delta\omega_{ES1}$ (ppm) | -2.36 ± 0.01 | -2.38 ± 0.01 |
| $R_1$ ($s^{-1}$) | 24.4 ± 0.3 | 22.4 ± 0.1 |
| $R_2$ ($s^{-1}$) | 41.1 ± 0.4 | 40.8 ± 0.4 |
| Red. $\chi^2$ | 12.9 | 12.4 |

| T = 10˚C | | |
|---|---|---|
| Parameter | 8OG-H7 900 MHz | 8OG-H7 900 MHz high spin-lock power |
| $pop_{ES1}$ (%) | 4.7 ± 2.6 | - |
| $pop_{ES2}$ (%) | - | 0.06 ± 0.04 |
| $k_{ex,ES1}$ ($s^{-1}$) | 22 ± 10 | - |
| $k_{ex,ES2}$ ($s^{-1}$) | - | 2790 ± 2313 |
| $\Delta\omega_{ES1}$ (ppm) | -2.27 ± 0.01 | - |
| $\Delta\omega_{ES2}$ (ppm) | - | -2.6 ± 0.1 |
| $R_1$ ($s^{-1}$) | 6.59 ± 0.01 | 6.78 ± 0.04 |
| $R_2$ ($s^{-1}$) | 30.8 ± 0.3 | 28.3 ± 0.2 |
| Red. $c^2$ | 11.0 | 12.7 |

**Supplementary Table 3.** Summary of exchange parameters obtained from fitting $^{13}C$ $R_{1\rho}$ data measured on labeled hp$^{8OG•A}$ A-C1' and A-C8 at T = 10˚C and pH 7.4 in 90% $H_2O$:10% $D_2O$. Red. $\chi^2$ denotes the reduced $\chi^2$ obtained on fitting the RD data. ES1 corresponds to 8OG$_{anti}$•A$_{anti}$ and ES2 corresponds to 8OG$_{anti}$•A$_{syn}$.

| T = 10˚C | | |
|---|---|---|
| Parameter | A-C8 | A-C1′ |
| $pop_{\mathrm{ES1}}$ (%) | 8.3 ± 0.9 | - |
| $pop_{ES2}$ (%) | - | 0.15 ± 0.04 |
| $k_{\mathrm{ex,ES1}}$ ($s^{-1}$) | 24 ± 4 | - |
| $k_{\mathrm{ex,ES2}}$ ($s^{-1}$) | - | 1473 ± 617 |
| $\Delta\omega_{\mathrm{ES1}}$ (ppm) | 1.6 ± 0.1 | - |
| $\Delta\omega_{\mathrm{ES2}}$ (ppm) | - | 3.5 ± 0.4 |
| $R_1$ ($s^{-1}$) | 2.9 ± 0.1 | 2.8 ± 0.1 |
| $R_2$ ($s^{-1}$) | 33.3 ± 0.2 | 21.2 ± 0.2 |
| Red. $\chi^2$ | 0.42 | 0.54 |

**Supplementary Table 4.** Summary of exchange parameters obtained from [1]H CEST on all constructs at T = 15°C where the residue exhibited exchange based on AIC/BIC model selection and the Δω was not the result of NOE effects.

| Construct | Parameter | T17 | T15 |
|---|---|---|---|
| $hp^{A\text{-}T}$ | $pop_{ES}$ (%) | 0.38 ± 0.01 | 0.06 ± 0.01 |
| | $k_{ex,ES}$ ($s^{-1}$) | 5298 ± 637 | 3186 ± 762 |
| | $\Delta\omega_{ES}$ (ppm) | -0.9 ± 0.1 | -1.9 ± 0.1 |
| | $R_1$ ($s^{-1}$) | 2.42 ± 0.01 | 5.68 ± 0.02 |
| | $R_2$ ($s^{-1}$) | 12.2 ± 0.4 | 15.1 ± 0.3 |
| | Red. $\chi^2$ | 389.9 | 311.8 |
| $hp^{G•A}$ | Parameter | T17 | G15 |
| | $pop_{ES}$ (%) | 0.4 ± 0.1 | 4.6 ± 0.1 |
| | $k_{ex,ES}$ ($s^{-1}$) | 1840 ± 125 | 1014 ± 24 |
| | $\Delta\omega_{ES}$ (ppm) | -0.94 ± 0.02 | -1.5 ± 0.01 |
| | $R_1$ ($s^{-1}$) | 2.00 ± 0.01 | 5.68 ± 0.01 |
| | $R_2$ ($s^{-1}$) | 11.2 ± 0.2 | 16.5 ± 0.2 |
| | Red. $\chi^2$ | 287 | 6.7 |
| $hp^{8OG•A}$ | Parameter | T17 | 8OG15 |
| | $pop_{ES}$ (%) | 0.63 ± 0.07 | 3 ± 1 |
| | $k_{ex,ES}$ ($s^{-1}$) | 4158 ± 456 | 62 ± 19 |
| | $\Delta\omega_{ES}$ (ppm) | -0.86 ± 0.04 | -2.31 ± 0.01 |
| | $R_1$ ($s^{-1}$) | 2.21 ± 0.01 | 7.8 ± 0.1 |
| | $R_2$ ($s^{-1}$) | 21.5 ± 0.3 | 30.6 ± 0.3 |
| | Red. $\chi^2$ | 17.3 | 7.6 |
